# Nothing else matters? Tree diameter and living status have more effects than biogeoclimatic context on microhabitat number and occurrence: an analysis in French forest reserves

**DOI:** 10.1101/335836

**Authors:** Yoan Paillet, Nicolas Debaive, Frédéric Archaux, Eugénie Cateau, Olivier Gilg, Eric Guilbert

## Abstract

Managing forests to preserve biodiversity requires a good knowledge not only of the factors driving its dynamics but also of the structural elements that actually support biodiversity. Tree-related microhabitats (e.g. cavities, cracks, conks of fungi) are tree-borne features that are reputed to support specific biodiversity for at least a part of species’ life cycles. While several studies have analysed the drivers of microhabitats number and occurrence at the tree scale, they remain limited to a few tree species located in relatively narrow biogeographical ranges. We used a nationwide database of forest reserves where microhabitats were inventoried on more than 22,000 trees. We analysed the effect of tree diameter and living status (alive or dead) on microhabitat number and occurrence per tree, taking into account biogeoclimatic variables and tree genus.

We confirmed that larger trees and dead trees bore more microhabitats than their smaller or living counterparts did; we extended these results to a wider range of tree genera and ecological conditions than those studied before. Contrary to our expectations, the total number of microhabitat types per tree barely varied with tree genus – though we did find slightly higher accumulation levels for broadleaves than for conifers – nor did it vary with elevation or soil pH, whatever the living status. We observed the same results for the occurrence of individual microhabitat types. However, accumulation levels with diameter and occurrence on dead trees were higher for microhabitats linked with wood decay processes (e.g. dead branches or woodpecker feeding holes) than for other, epixylic, microhabitats such as epiphytes (ivy, mosses and lichens).

Promoting large living and dead trees of several tree species may be a relevant, and nearly universal, way to favour microhabitats and enhance the substrates needed to support specific biodiversity. In the future, a better understanding of microhabitat drivers and dynamics at the tree scale may help to better define their role as biodiversity indicators for large-scale monitoring.

## Introduction

Small natural features are structural habitat elements that have a disproportionately important role for biodiversity related to their actual size [1]. Taking these features into account in biodiversity conservation strategies is a crucial step in science-based decision making [2]. Identifying such structural features in a tri-dimensional forest environment is quite challenging since their number and variety is potentially infinite. Small natural features include, for example, large old trees [3] as well as tree-borne structures. While large old trees are disappearing at the global scale [4], their importance for biodiversity has not yet been fully elucidated, not to mention the peculiar structures they may bear (eg. cracks, cavities, epiphytes), also known as ‘tree-related microhabitats’ (hereafter ‘microhabitats’ [5]). Microhabitats have recently aroused the interest of scientists and forest managers alike since these structures can be a substrate for specific forest biodiversity [6], and can ultimately serve as forest biodiversity indicators [5, 7, 8]. Their conservation has hence become an issue in day-to-day forest management, as have large old trees and deadwood [9, 10]. However, our understanding of the drivers and dynamics influencing these microhabitats, notably at the tree scale, remains incomplete [11]. Tree diameter and living status (living vs. dead trees) are key factors for microhabitat diversity at the tree scale [12–14]. Larger trees are likely to bear more microhabitats than smaller ones, as they have experienced more damage, injuries and microhabitat-creating events (e.g. woodpecker excavation, storms, snowfalls). Similarly, gradually decomposing dead trees are likely to bear more microhabitats than living trees and play a role as habitat and food sources for many microhabitat-creating species [15]. Nevertheless, the relationships between microhabitats and tree characteristics have only been demonstrated on a limited number of tree species involving at most a few thousand observations at the tree level (e.g. [11–13]), which have been carried out within a limited biogeographical range (e.g. in Mediterranean forests [16], the French Pyrenees [12] or in Germany [17, 18]). Consequently, it remains to be understood whether the observed relationships between tree characteristics and microhabitats – even though they seem to be relatively consistent across studies – are merely idiosyncratic, notably in terms of magnitude. Large databases making larger-scale analyses possible are rare (but see [19]), mainly due to a lack of homogeneity in the typologies used to inventory microhabitats [5] and a lack of forest microhabitat monitoring initiatives. Large-scale data are, nonetheless, crucial to better understanding the potential variations in the relationships between microhabitat and biotic (e.g. tree species) or abiotic (e.g. climatic) factors, with a view to validating microhabitats as potential biodiversity indicators at various scales [7, 8, 18].

We used a nationwide database resulting from standardized monitoring in forest reserves, where microhabitats have been inventoried since 2005. We analysed the influence of individual tree diameter and living status on the number and occurrence of microhabitat types at the tree level. We expected the number and occurrence of microhabitats per tree to increase with diameter and to be higher on dead than on living trees. We assessed the influence of tree species and biogeoclimatic variables on these relationships, expecting that microhabitat dynamics (or accumulation rate per tree) would be tree-species dependent and would vary with abiotic context (higher accumulation rates in harsher conditions: e.g. at high elevations or on acidic soils). Ultimately, the aim of this study was to provide forest managers with a better science-based knowledge of microhabitats in the forest ecosystem, thus allowing them to adapt their management to specific local contexts.

## Materials and methods

### Database structure

We worked with a nationwide database compiled from a monitoring program in French forest reserves. Since 2005, a systematic permanent plot network has gradually been set-up on a voluntary basis in forest reserves. The main objectives of this network are (i) to better understand the dynamics of forest ecosystems subjected to varying degrees of management, (ii) to provide reserve managers with quantitative data on the flux of living and dead trees at the site scale, and (iii) to ultimately provide guidelines for establishing management plans. The full database currently includes 107 reserves for a total of 8190 plots (83180 living and 19615 dead trees, snags or stumps). The forest reserves in the database actually encompass three broad types of protection status. First, (i) strict forest reserves, where harvesting has been abandoned for a variable timespan and (ii) special forest reserves, where management is targeted towards specific biodiversity conservation measures (e.g. preservation of ponds). These two types are owned and managed by the French National Forest Service. The third type, nature reserves, on the other hand, where management varies from abandonment to classic wood production, may be of various ownership types (state, local authorities, private). It should be noted that no homogeneous data on management intensity or time since last harvesting could be gathered at the plot level for all the reserves in the database. However, Vuidot et al. [13] showed that management has a limited effect on microhabitat number and occurrence at the tree level. We thus assumed that management differences would not play a significant role at the tree scale and therefore, did not take management type or intensity into account in our analyses (but see discussion).

### Stand structure and microhabitat inventories

On each plot, we combined two sampling methods to characterise forest stand structure [20]. For all living trees with a diameter at breast height (DBH) above 30 cm, we used a fixed angle plot method to select the individuals comprised within a relascopic angle of 3%. Practically, this meant that sampling distance was proportional to the apparent DBH of a tree. For example, a tree with a DBH of 60 cm was included in the sample if it was within 20 m of the centre of the plot. This particular technique allowed us to better account for larger trees at a small scale. All other variables were measured on fixed-area plots. Within a fixed 10-m (314 m^2^) radius around the plot centre, we measured the diameter of all living trees and snags (standing dead trees with a height > 1.30 m) from 7.5 to 30 cm DBH. Within a 20-m radius (1256 m^2^), we recorded all snags with a diameter > 30 cm. Whenever possible, we identified all trees, both alive and dead, to species level. In the subsequent analyses, we grouped some tree species at the genus level to have sufficient representation in terms of tree numbers. This resulted in the following groups: ash (Fraxinus excelsior L.), beech (Fagus sylvatica L.), chestnut (Castanea sativa Mill.), fir (Abies alba Mill.), hornbeam (Carpinus betulus L.), larch (Larix decidua Mill.), maple (90% sycamore maple, Acer pseudoplanatus L.), oak (80% sessile, Quercus petraea (Matt.) LIebl., and pedunculate, Q. robur L., oaks combined, 15% oaks identified to the genus level only, 5% other oaks – mainly Mediterranean), pine (64% Scots pine, Pinus sylvestris L., 22% mountain pine, Pinus mugo Turra), poplar (Populus spp.) and spruce (Picea abies (L.) H. Karst. We assumed that tree genus, rather than species, influenced the relationships we were studying. Unidentified species were excluded from the analyses.

We visually inspected all selected standing trees for microhabitats and recorded their presence on each tree. Observers attended a training session and were given a field guide with pictures to help them better determine microhabitat types and detailed criteria to include in the inventories. Although inventory methods have recently improved [5, 21], we assumed that the method we used limited any potential observer effect linked with microhabitat inventories [22].

Different microhabitat typologies were used concomitantly during the inventories and harmonization has been lacking since 2005. Consequently, we only retained data with a homogeneous typology. We preferred this solution rather than grouping microhabitat types to avoid coarser classification with too much degradation of the original dataset.

### Data selection and biogeoclimatic variables extraction

First, we focused on the microhabitat typology that was used for the largest number of plots and sites (Table 1). This reduced the dataset to 43 sites comprising 3165 plots (Figure 1, Table S1). Second, the smallest trees (7.5 ≤ DBH ≤17.5 cm) accounted for 36% of the trees in the database but were also the least likely to bear microhabitats [12, 13]. We therefore excluded this category from the dataset to avoid zero-inflation in the subsequent models. Third, previous studies had shown that tree living status (i.e. living vs. dead trees, see below) is a major driver of microhabitat occurrence and density [12, 13]. To properly account for this variable in our statistical models, we excluded all tree species/genera with less than 50 standing dead trees or snags in the dataset (ie. ash, chestnut, hornbeam, larch, maple, poplar, see Table 2 for distribution by genus and diameter classes and Supplementary Material, Figure S1, for a calculation based on a larger subset of living trees). The final dataset comprised 2783 plots distributed over 43 sites, for a total of 22307 trees (20312 living and 1995 dead trees belonging to five genera of both dead and living trees, Table 2).

**Figure 1:**
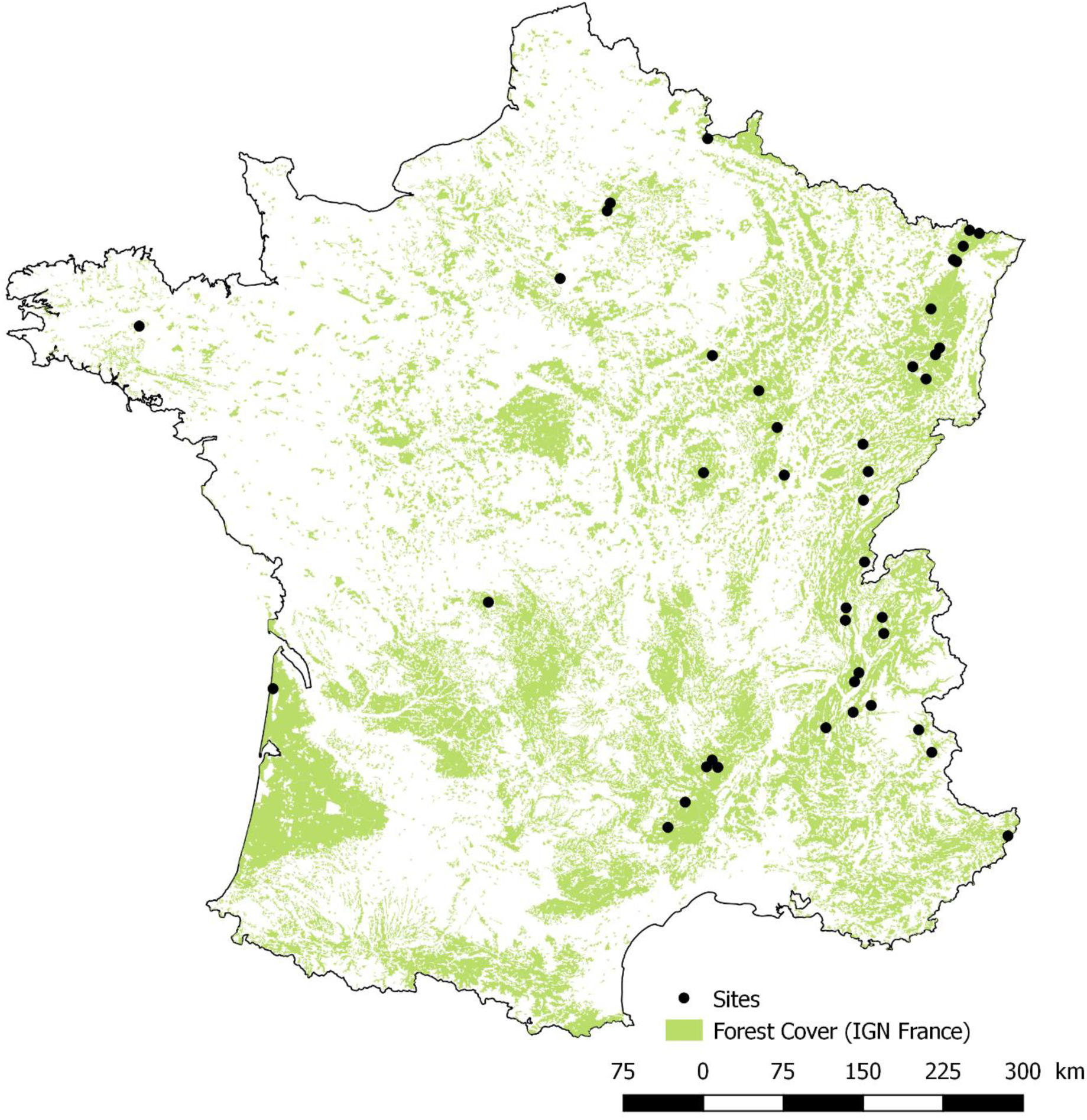
Location of the study sites

**Table 1:**
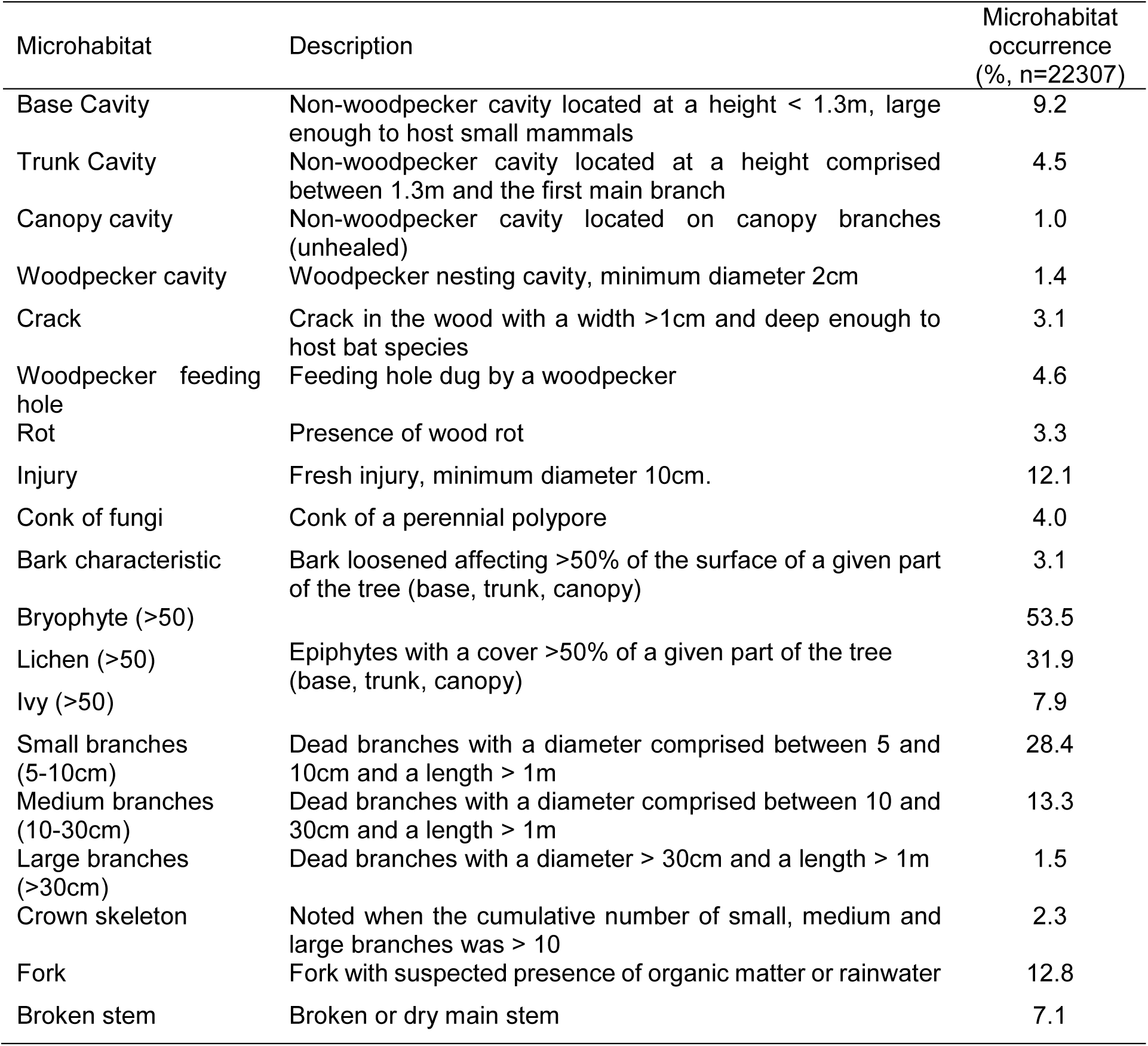
Microhabitat typology

**Table 2:**
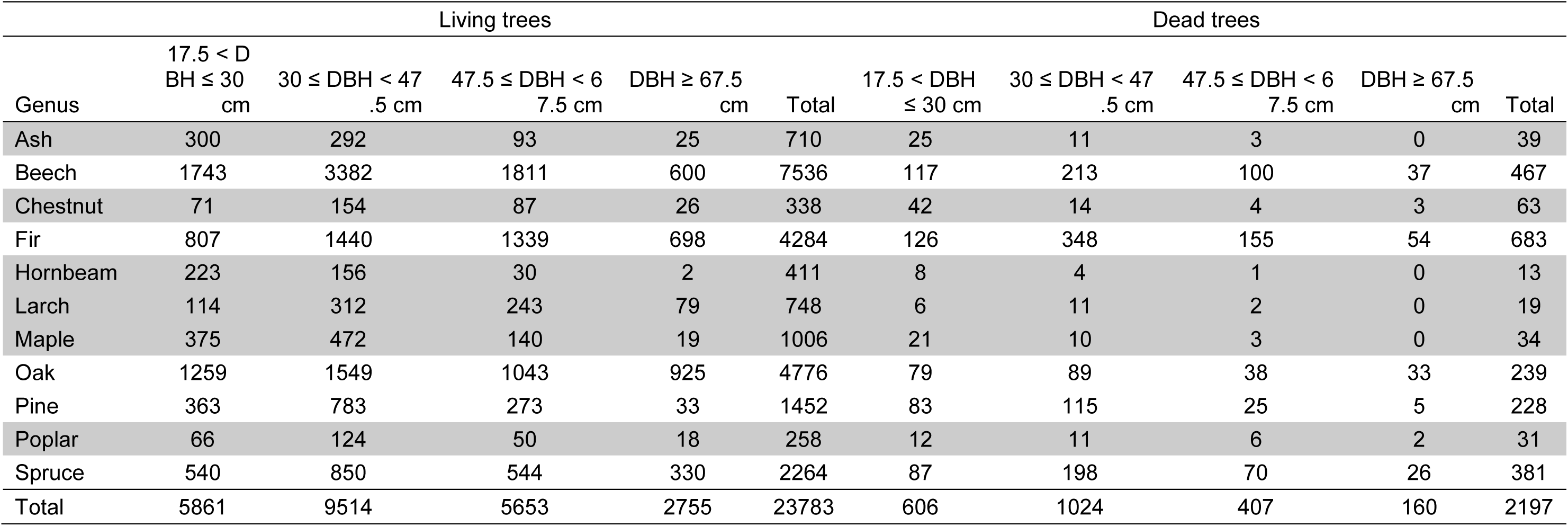
Distribution of the data by genus and Diameter at Breast Height (DBH) classes. Genera in grey were excluded from the main analyses due to an insufficient number of occurrences of dead trees; in this case, only living trees were analysed (see Supplementary Materials: Figure S1). ash: Fraxinus excelsior; beech: Fagus sylvatica; chestnut: Castanea sativa; fir: Abies alba; hornbeam: Carpinus betulus; larch: Larix decidua; maple: Acer spp., oak: Quercus spp.; pine: Pinus spp.; poplar: Populus spp.; and spruce: Picea abies.

In addition, we gathered different biogeoclimatic data from various sources to reflect plot characteristics:

- annual mean temperature (bio1) and precipitation (bio12) from the Worldclim2 database [23];
- elevation, aspect and slope from the national digital elevation model (resolution 30 m);
- soil plant-bioindicated pH from the National Forest Inventory [24].

### Statistical analyses

Following Zuur et al. [25], preliminary data exploration did not reveal any potential variation in the relationship between microhabitat metrics and any of the biogeoclimatic variables mentioned above, apart from pH and elevation. We therefore kept pH and elevation only in the analyses described below. However, elevation correlated strongly to tree species; indeed, only beech and pine were distributed over the whole elevation gradient while the other species were elevation-dependent. Conversely, genera were relatively well distributed over the pH gradient.

We used DBH, living status (alive vs. dead) and genus (beech, fir, oak, pine and spruce) as explanatory variables and included second and third order interactions between DBH, living status and genus in the models. We added elevation and pH as covariables, but only included pH in the second order interactions. Since beech and pine were not strongly biased by elevation, we added elevation in the second order interactions for these two genera in two separate analyses.

To model the total number of microhabitat types per tree, we used generalised linear mixed models (GLMMs, library glmmTMB, [26]) with a Poisson error distribution for count data and plot identity nested within site as a random variable. We also modelled the occurrence of each microhabitat type, but with a binomial error distribution for binary data. We tested differences in microhabitat numbers and occurrences between living and dead trees with post-hoc multi-comparison Tukey tests for a fixed mean DBH (44 cm; function cld, library emmeans [27]). Dispersion diagnostics revealed under-dispersed model estimations, which may cause a type II error rate inflation [28]. However, since there was no simple way to account for that in a frequentist framework, we kept the results while bearing in mind that they were undoubtedly conservative despite the large number of observations we analysed. In addition, we focused our interpretations on the magnitude of the results rather than their statistical significance (see e.g. [29]). We processed all the analyses with the R software v. 3.4.3 [30].

## Results

### Number of microhabitat types per tree

Estimates for all single parameters were significant in the model, except for soil pH, while second and third order interactions were less often significant (see Supplementary Materials, Table S2). All tree genera but pine had higher microhabitat numbers on dead than on living trees. Overall, the difference was the highest for oak (22% more microhabitats on dead than on living trees, for a mean DBH of 44 cm, Table 3); the other genera had around 10-15% more microhabitats on dead than on living trees. Globally, the number of microhabitats per tree increased with tree diameter, both for living and dead trees (Figure 2). However, the accumulation of microhabitats with diameter varied with genus (the two broadleaves’ genera investigated, beech and oak, had higher accumulation levels than the three conifers’ genera, fir, pine, spruce), and according to living status (dead versus living trees, except for pine; Figure 2, Supplementary Materials, Table S2). These results were generally consistent with those obtained with the analyses concerning a higher number of genera but for living trees only (Figure S1). Broadleaves (ash, beech, chestnut, hornbeam, maple, oak, poplar) showed higher microhabitat accumulation rates than conifers (fir, larch and spruce). Only pine showed accumulation rates comparable to broadleaves (Figure S1).

**Table 3:** Percentage of difference in number of microhabitats between living and dead trees for a mean Diameter at Breast Height (DBH = 44 cm) calculated as [(Microhabitats dead trees – Microhabitats living trees) / (Microhabitats dead trees + Microhabitats living trees)] x 100. An * indicates a significant (p<0.05) difference based on post-hoc Tukey tests for a mean DBH. Values close to −100 correspond to cases where microhabitats were quasi-absent on dead trees (resp. 100 for living trees). Figures in brackets are absolute values for dead and living trees respectively. Beech: Fagus sylvatica; fir: Abies alba; oak: Quercus spp.; pine: Pinus spp.; and spruce: Picea abies.

| Microhabitats | Beech | Fir | Oak | Pine | Spruce |
| --- | --- | --- | --- | --- | --- |
| All | 14.9*<br>[2.27-1.681] | 12.7*<br>[1.649-1.278] | 21.5*<br>[2.789-1.804] | 14.8<br>[1.511-1.121] | 11.4*<br>[1.464-1.164] |
| Base cavities | 18.6<br>[0.057-0.039] | 29.7<br>[0.022-0.012] | 4.6<br>[0.02-0.018] | 61.3<br>[0.008-0.002] | -32.9<br>[0.015-0.03] |
| Trunk cavities | 41.4*<br>[0.072-0.03] | 71.4<br>[0.02-0.003] | 49.3*<br>[0.049-0.017] | 79.9*<br>[0.021-0.002] | 86.8*<br>[0.01-0.001] |
| Canopy cavities | -44.9<br>[0.001-0.001] | -10.5<br>[<0.001-<0.001] | 10.0<br>[0.002-0.002] | -100<br>[<0.001-0.001] | 100<br>[<0.001-<0.001] |
| Woodpecker cavities | 77.9*<br>[0.029-0.004] | 39.7<br>[0.006-0.002] | 64.6*<br>[0.018-0.004] | 63.7<br>[0.015-0.003] | 26.3<br>[0.003-0.002] |
| Cracks | 42.9*<br>[0.035-0.014] | 41.4<br>[0.01-0.004] | 82.8*<br>[0.039-0.004] | -66.2<br>[0.001-0.004] | 54.4<br>[0.016-0.005] |
| Woodpecker feeding holes | 97.5*<br>[0.285-0.004] | 98.6*<br>[0.362-0.003] | 95.8*<br>[0.362-0.008] | 95.6*<br>[0.13-0.003] | 97.9*<br>[0.184-0.002] |
| Rot | 45.9*<br>[0.039-0.014] | 22.3<br>[0.013-0.008] | 90.3*<br>[0.138-0.007] | 82.2*<br>[0.013-0.001] | 80.3*<br>[0.027-0.003] |
| Injuries | -67.4*<br>[0.015-0.075] | -82.8*<br>[0.006-0.06] | -62.5*<br>[0.011-0.049] | -74.5*<br>[0.004-0.028] | -89.3*<br>[0.005-0.086] |
| Conks of fungi | 96.1*<br>[0.37-0.007] | 98.0*<br>[0.271-0.003] | 86.9*<br>[0.076-0.005] | 94.1*<br>[0.062-0.002] | 96.2*<br>[0.151-0.003] |
| Bark characteristics | 92.1*<br>[0.061-0.003] | 94.0*<br>[0.049-0.002] | 98.6*<br>[0.262-0.002] | 96.9*<br>[0.056-0.001] | 98.5*<br>[0.106-0.001] |
| Moss cover >50% | -18.1*<br>[0.458-0.66] | -37.7*<br>[0.154-0.341] | -56.6*<br>[0.225-0.809] | 55.0<br>[0.105-0.03] | 6.0<br>[0.092-0.082] |
| Lichen cover >50% | -61.1*<br>[0.029-0.121] | -71.9*<br>[0.035-0.216] | -29.1<br>[0.074-0.135] | -32.7<br>[0.04-0.08] | -75.8*<br>[0.011-0.081] |
| Ivy cover >50% | -25.6<br>[0.001-0.002] | -54.2<br>[<0.001-0.002] | -4.5<br>[0.003-0.004] | 25.5<br>[0.002-0.001] | -30.9<br>[0.002-0.003] |
| Small branches | -82.7*<br>[0.015-0.153] | -52.8*<br>[0.031-0.1] | -88.1*<br>[0.02-0.318] | -84.6*<br>[0.031-0.371] | -46.7<br>[0.02-0.056] |
| Medium branches | -58.8*<br>[0.012-0.045] | 81.7*<br>[0.052-0.005] | -59.5*<br>[0.043-0.17] | -48.9<br>[0.037-0.106] | -39.4<br>[0.002-0.004] |
| Large branches | 33.7<br>[0.003-0.001] | 42.8<br>[<0.001-<0.001] | -52.8<br>[0.002-0.006] | 54.2<br>[0.008-0.002] | -100<br>[<0.001-<0.001] |
| Crown skeleton | 98.3*<br>[0.003-<0.001] | 74.6<br>[<0.001-<0.001] | 97.4*<br>[0.003-<0.001] | 85.3*<br>[0.006-<0.001] | 91.2*<br>[0.017-0.001] |
| Forks | -94.3*<br>[0.002-0.075] | -72.4*<br>[0.003-0.021] | -48.9<br>[0.02-0.059] | -82.9<br>[0.003-0.032] | -67.9*<br>[0.004-0.019] |
| Broken stem | 12.1<br>[0.029-0.023] | 0.4<br>[0.033-0.032] | -1.6<br>[0.021-0.021] | -38.8<br>[0.012-0.028] | -9.8<br>[0.021-0.026] |

**Figure 2:**
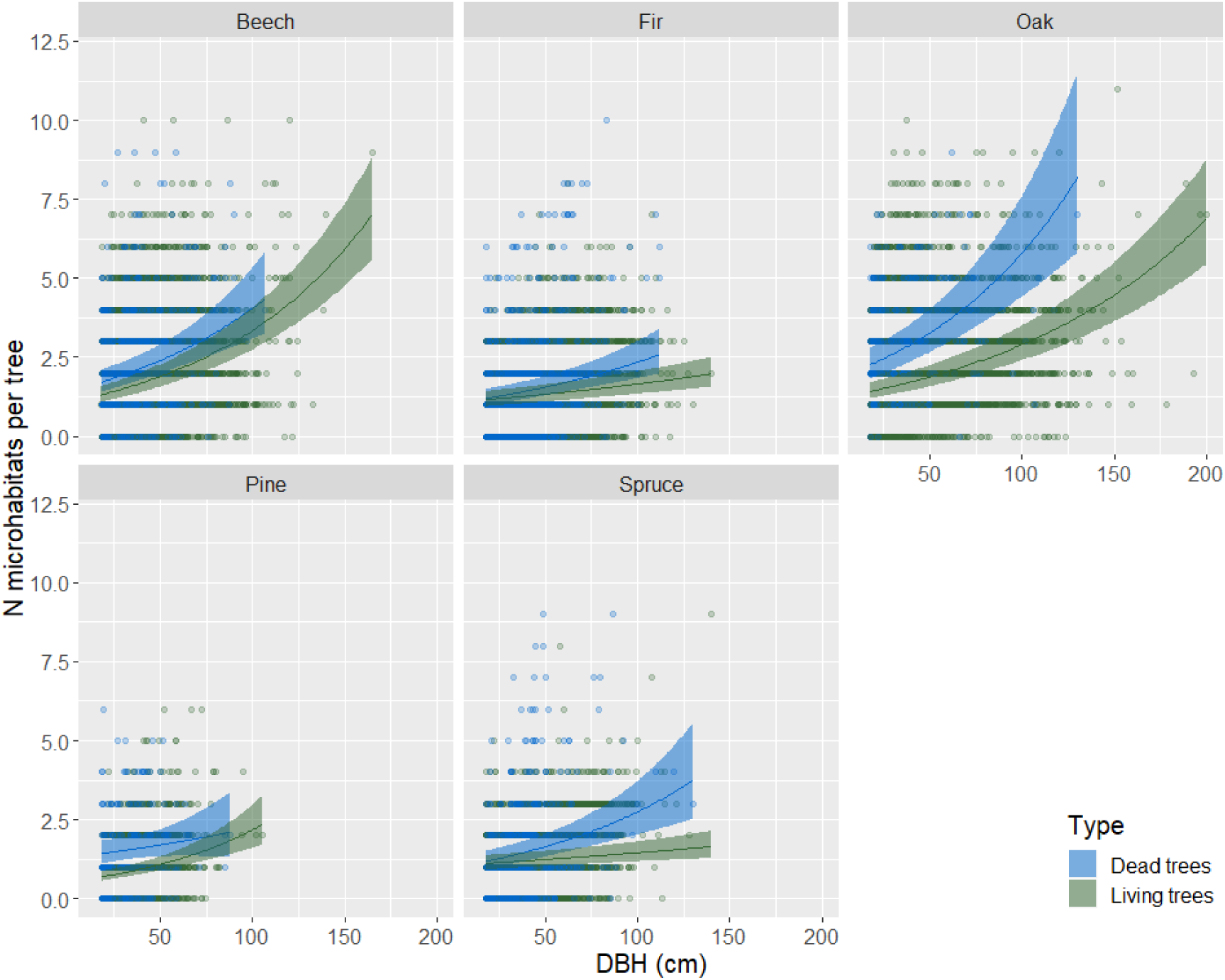
Relationship between number of microhabitats (N microhabitats per tree) and Diameter at Breast Height (DBH, cm) by genera (beech: Fagus sylvatica; fir: Abies alba; oak: Quercus spp., pine: Pinus spp. and spruce: Picea abies) and living status (living vs. dead standing trees). Lines represent estimates from generalized mixed effect models with a Poisson error distribution and plot nested in site as a random effect. Ribbons show the 95% confidence intervals of the mean. For this representation, pH and elevation were held constant (mean values in our data set).

Number of microhabitats increased significantly with elevation, but not with soil pH. However, higher soil pH had a positive effect on the accumulation of microhabitats with DBH (the second order interaction was significant), mostly on dead trees (Supplementary Materials, Table S2). Still, the effects of elevation and soil pH remained small compared to those of DBH and living status.

For beech and pine, the overall results converged with those of the complete model. Soil pH and elevation only had significant effects in the interaction terms (Supplementary Materials: Table S4): increasing soil pH increased microhabitat accumulation with DBH for both species, with a stronger effect for pine than for beech. On the other hand, increasing soil pH decreased microhabitat richness on living compared to dead trees. Elevation interacted significantly with living status for beech only, and almost doubled the difference between living and dead trees, whereas for pine, the effects were only marginally significant (p<0.1), though high in magnitude.

### Occurrence of microhabitat types per tree

Six microhabitats out of twenty generally occurred more frequently on standing deadwood than on living trees, though this was not systematic for all genera or even for living status: trunk cavities (except fir), woodpecker feeding holes (Figure 3), rot (except fir), conks of fungi, bark characteristics and crown skeleton (except fir, Table 3 and Supplementary Materials, Table S5). We observed the strongest differences for woodpecker feeding holes: whatever the genera, they virtually only occurred on standing dead trees (i.e. they were nearly absent from living trees, Figure 3, Table 3). Conversely, injuries, dead branches whatever their size and forks (broadleaves only) occurred more frequently on living trees. Magnitudes for microhabitats more frequent on living trees were around 60% to 90% (Table 3).

**Figure 3:**
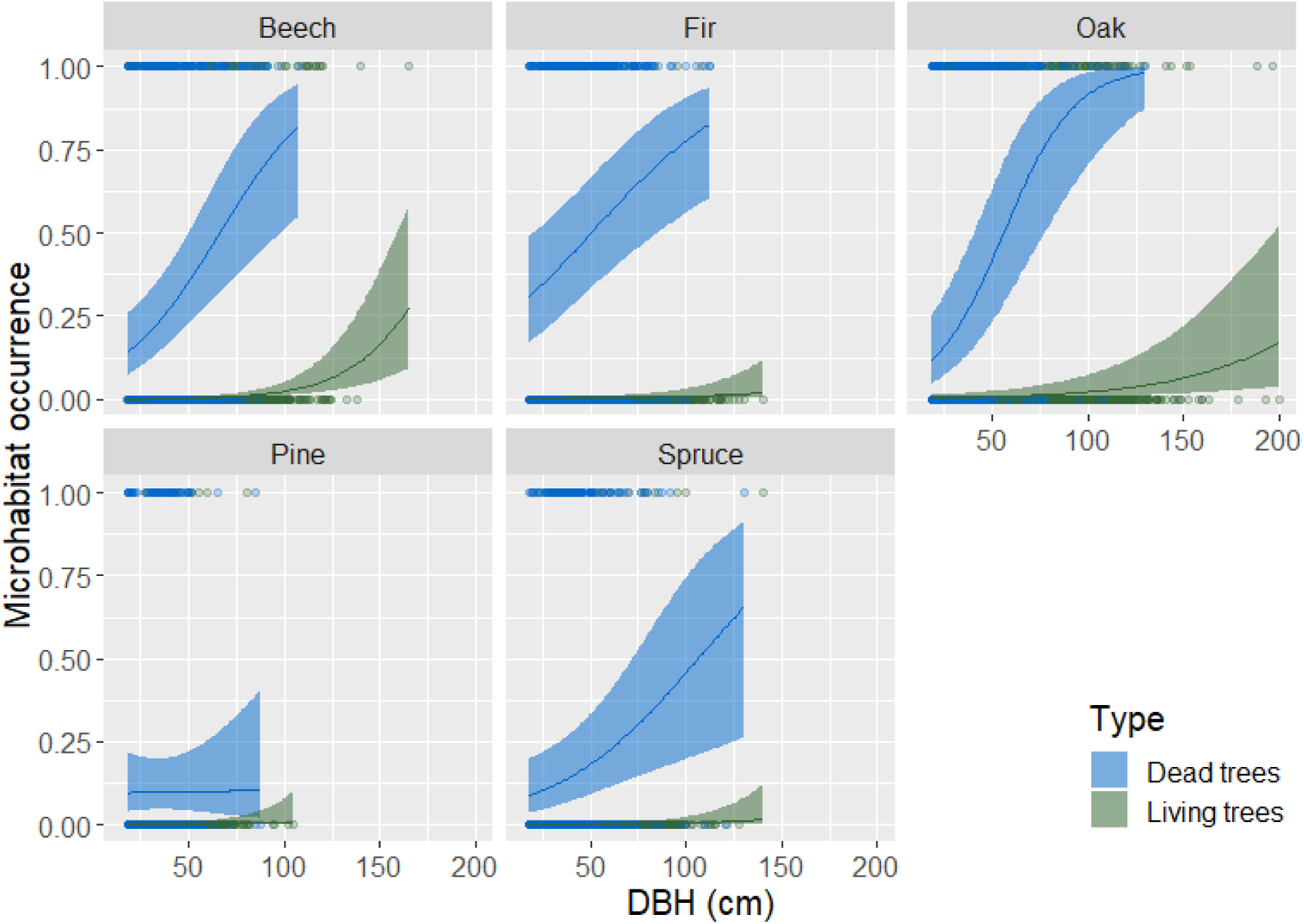
Relationship between occurrence of woodpecker feeding holes and Diameter at Breast Height (DBH, cm) by genera (beech: Fagus sylvatica; fir: Abies alba; oak: Quercus spp., pine: Pinus spp. and spruce: Picea abies) and living status (living vs. dead standing trees). Lines represent estimates from generalized mixed effect models with a binomial error distribution. Ribbons show the 95% confidence interval of the mean. For this representation, pH and elevation were held constant. See Supplementary materials, Figure S2, for all microhabitat types.

For most microhabitats, the probability of occurrence increased with DBH both for living and dead trees, with the remarkable exceptions of canopy cavities, woodpecker cavities and crown skeletons (Supplementary Materials: Figure S2, Table S5). However, the magnitude of the relation varied with tree genus and living status. For some microhabitat types, the increase in probability of occurrence with DBH was stronger for dead than for living trees, e.g.: +35% base and trunk cavities on dead vs. +18% on living beech; +23 to +42% for woodpecker feeding holes on dead vs. +0.2 to +3% on living trees (Table S3). Conversely, the increase in probability of occurrence of small and medium dead branches was stronger for living trees (e.g. +53% medium dead branches on living vs. 0.7% on dead oak) and, to a lesser extent, for mosses on beech and fir (+20% and +24% on living trees, vs. +9% and +16% on dead trees, respectively). All other increments with DBH for living trees were smaller, generally below 10%. Note that in some cases, due to the very limited number of occurrences for some microhabitats on certain tree genera, the estimates proved unreliable (huge confidence intervals, e.g. canopy cavities on oak, pine and spruce, Supplementary Materials: Figure S2, Table S5).

Elevation had an overall negative effect on microhabitat occurrence, except for trunk cavities, lichens and forks. Conversely, soil pH tended to have a positive effect on microhabitat occurrence, except for conks of fungi. More interestingly, increasing soil pH had a positive effect on the accumulation of some microhabitats when coupled with DBH (indicated by a significant interaction term), but a negative effect on occurrence on living trees (Supplementary Materials: Table S5). All these significant effects exhibited widely varying levels of magnitude, and in several cases, the estimates were rather imprecise (Supplementary Materials: Figure S2, Table S5).

## Discussion

Numerous recent studies in a variety of contexts have shown that the number of microhabitats per tree as well as the occurrence of some microhabitat types increase with tree diameter [11, 14, 16]; these studies also evidenced higher occurrence levels on dead than on living trees [12, 13]. Our nationwide study based on a large database confirmed these relationships and extended them to a larger range of tree genera under wider biogeographical conditions. Indeed, our results include five tree genera for both living and dead trees and eleven genera when only living trees were considered (Supplementary Materials: Figure S1).

### Dead trees bear more microhabitats than living trees

Standing dead trees contribute significantly to the supply of microhabitats; overall, they bore 10 to 20% more microhabitats than their living counterparts in our dataset comprising five genera. Dead trees often bear considerably more microhabitats than living trees when individual microhabitat types are analysed (e.g. woodpecker feeding holes – Figure 3 – or bark characteristics). Once dead, standing trees are affected by decomposition processes that trigger microhabitat genesis [15]. Standing dead trees also constitute privileged foraging grounds for a number of species [5, 7, 8], including woodpeckers [31, 32]. In particular, insect larvae or ants that live under the bark of more or less recently dead trees provide a non-negligible part of some birds’ diet [8, 33, 34]. Furthermore, as living trees also bear microhabitats, it seems logical that many of these would persist when the tree dies and would continue to evolve, or possibly even condition the presence of other microhabitats linked with the decaying process [15]. For example, injuries caused by logging, branch break or treefall could begin to rot and then slowly evolve into decay cavities [5, 35]. These successional changes are likely to explain why these microhabitats types are more numerous on dead trees. The only exceptions to this global pattern concerned epiphytes and forks with accumulated organic matter, which both tend to be more numerous on living trees. Ivy, mosses and lichens are likely to benefit from bark characteristics (e.g. pH, [36]) occurring only on living trees. Epiphytes, especially slow-growing mosses and lichens, require a relatively stable substrate to take root and develop [37]. Stability is lost when bark loosens and falls off during tree senescence, and this could cause epiphytic abundance to decrease. In a nutshell, decaying processes linked to the tree’s death reveal a clear difference between microhabitats that are linked to decay (i.e. saproxylic microhabitats, sensu [5]) and those that are not – or less so (i.e. epixylic microhabitats).

Nearly all previous studies comparing microhabitat numbers on living and dead trees found more microhabitats on dead trees (see [17]). However, the difference varies across studies, from 1.2 times as many microhabitats in Mediterranean forests [16] and twice as many in five French forests [13] to four times as many on habitat trees in south-western Germany [38]. Our results ranged from 1.1 to 1.2 times as many microhabitats on dead as on living trees, which is of a slightly lower order of magnitude than previously reported. This surprising result may be due to the fact that our study encompassed more species with a lower microhabitat bearing potential (namely conifers). Yet, even for the same species analysed in previous studies (e.g. beech), the levels we observed were lower. Since we found only small effects of pH and elevation, this finding seems to indicate that the difference in magnitude is not due to biogeographical variation.

### Number and occurrence of microhabitats increase with tree diameter

We confirmed that both microhabitat number and occurrence increase with tree diameter but, contrary to expectations ([11–13], but see [14]), tree genus had a limited effect on this relationship, with only slightly higher microhabitat accumulation levels on broadleaves than on conifers. Almost all microhabitat types taken individually showed the same increasing trends with tree DBH, but there were considerable variations in magnitude. Larger (living) trees have generally lived longer than smaller ones, and are consequently more likely to have suffered more damage during their lifespan due to meteorological events (storms, snowfall), natural hazards (rockfalls) or use by different tree- and wood-dependent species (woodpeckers, beetles, fungi, see e.g. [13, 39]). In some studies, doubling tree diameter (from 50 to 100 cm) has been shown to roughly double the number of tree microhabitats [13, 17, 18], though some studies have found multiples of up to four [38] or even five times [12] in certain cases. Again, our results showed magnitudes below the lower end of this range (the multiplication coefficient ranged from 1.2 to 1.4). This may be because the largest trees in our dataset were undoubtedly younger than those in the other studies, especially in studies on near-natural or long-abandoned forests [12, 13]. Indeed, since most of our sites had been (more or less) recently managed, selective felling may have cause trees with a given diameter to be younger than their counterparts in primeval forests, where competition levels may be higher and cause slower growth rates. At the individual microhabitat scale, dead branches were more likely to occur on large trees than on smaller trees; although this result seems quite obvious, it had rarely been quantified before. Larger trees have more, but also larger, branches likely to die from competition with neighbours, especially in broadleaves [40]. Indeed, oak and beech were the genera that showed the highest large dead branch accumulation rates with diameter in our analyses, while conifers had almost no large dead branches.

Cavity birds and bats are reputed to prefer larger trees for nesting or roosting [41, 42], since thicker wood surrounding the cavity provides a better buffered and more stable microclimatic conditions [43]. However, we did not confirm this relationship; the accumulation rates of woodpecker cavities with tree diameter were very weak and non-significant. The supposed relationship between tree diameter and woodpecker cavity occurrence seems hard to prove in the context of temperate European forests, at least with data from censuses comparable to ours (see [13] at the tree scale, or [44] at the stand scale); more targeted research focusing on this specific relationship is probably needed [31, 45]. Our results could also be linked to the non-linear dynamics [11] of this particular microhabitat. Some cavities in living beech can close back up when they are no longer used [pers. obs. Y.P.], and trees weakened by cavity digging can break, e.g. [45]. Other microhabitats, for instance conks of fungi, may also show non-linear dynamics linked with specific phenology [46]. In our study, the number and occurrence of microhabitats also increased with diameter in standing dead trees, sometimes at a higher rate than for living trees. The longer persistence of large dead trees compared to smaller ones [47] may combine the effects of increased damage due to hazards and the natural decaying processes described above. This probably explains the higher accumulation levels we observed in many cases, especially for saproxylic microhabitats (e.g. rot, feeding holes, trunk cavities). Once again, the only exception to this rule was the epiphytes: their probability of occurrence tended to increase with tree diameter but very noisily, both for living and dead trees. For such epiphytic organisms (ivy, mosses and lichens), larger scale processes and biogeoclimatic context (e.g. soil fertility, precipitation) is probably more important than individual tree characteristics [48]. This is suggested by the significant and rather strong effects of pH and elevation in our analyses (Supplementary Materials, Table S4).

### Limitations and research perspectives

Contrary to our expectations, we found a limited effect of biogeoclimatic variables on the relationship between microhabitats, tree diameter and living status. However, some specific interactions may exist, especially in the case of epiphytes [48], but that could not be evidenced by our approach. In addition, it was rather difficult to disentangle the effects of tree genus from those of the biogeoclimatic variables, since the distribution of most tree genera is driven largely by climate – apart from beech, and more marginally pine, which occur over broad bioclimatic gradients. However, even when we analysed beech and pine separately, we did not find any effect of soil pH or elevation on the number of microhabitats, and only slight effects on accumulation levels with diameter. These results need to be confirmed by further analyses with larger and more carefully controlled biogeographical gradients.

Our data from forest reserves potentially reflect a larger anthropogenic gradient than classical managed forests. Some of the reserves had not been harvested for several decades and exhibited characteristics of over-mature forests (see e.g. [20], who analysed some of the reserves included in this paper). On the other hand, their overall structure reflected relatively recent management abandonment – if any – since the reserves were marked by probable intensive use or previous harvesting over the past centuries, as is characteristic of western European forests [49]. This is testified to in the dataset we analysed by the relatively rare occurrence of dead standing trees, in particular those with a large diameter: standing dead trees represented a mere 10% of the total dataset and very large individuals (DBH > 67.5cm) only 1% (Table 2). As a consequence, despite the fact that we worked on an extended management gradient ranging from managed forests to unmanaged strict reserves, some of the elements characteristic of old-growth and over-mature forests were still lacking, especially large dead trees [50]. This truncated the relationships for the investigated set of microhabitats and made them imprecise for the larger diameter categories. Further research on the last remnant of old-growth primeval forests in Europe [51, 52] is therefore needed to bridge this gap and better understand microhabitat dynamics over the whole lifespan of the tree.

Compared to recent developments [5, 21], the microhabitat typology we used (Table 1) seems rather coarse or imprecise. This may explain why we were not able to confirm some of the effects mentioned in the literature; different microhabitats from a given group may have different requirements and dynamics (e.g. cavities dug by the black woodpecker vs. other woodpecker species). On the other hand, our descriptions allowed us to have enough occurrences in each type to analyse the combined effects of diameter and genus for almost all the microhabitat types in the typology. Our approach can be viewed as a compromise between providing the necessary sample size for statistical analyses and the degree of refinement in typology. The current developments mentioned above [5] will certainly help to homogenize data in the near future and to build larger, shared databases on common, comparable grounds.

Despite a training session prior to the inventories, observer effects cannot be totally ruled-out. Our censuses were mostly performed by non-specialists [22], contrary to the scientific studies previously published, and this may have led to the relatively low magnitudes observed, with the hypothesis that detection error is higher on one status (either dead or living trees) or one type of tree (e.g. small trees, which can be overlooked to the benefit of larger individuals). Such issues remain to be explored.

Finally, our models assumed – unrealistically as it turns out – that microhabitat number would increase exponentially with diameter. In fact, recent studies, as well as ecological theory (e.g. species-area relationship), tend to show a saturated (e.g. logarithmic or sigmoid) relationship between microhabitats and diameter. Models allowing for different link functions – probably within a Bayesian framework – will need to be tested to see whether they perform better than the ones used here (see e.g. [11]).

### Implications for forest management and biodiversity conservation

Large old trees are considered keystone small natural features in forest and agro-pastoral landscapes because of their disproportionate importance for biodiversity relative to their size [3]. This role for biodiversity is further enhanced by the ‘smaller’ natural features – microhabitats – they bear [7]. In our large-scale analysis, we confirmed and extended results previously observed only locally: most microhabitats occur on large trees, and even more on dead ones than on living ones. This relationship seems true for several tree genera included in this analysis, and across a large gradient of ecological conditions, with minor variations in accumulation rates with soil pH and elevation. As a consequence, conserving and recruiting large living and dead trees in daily forest management will enhance structural heterogeneity at the stand scale [6, 53], and favour a variety of tree-borne microhabitats, which could further help to better conserve specific forest biodiversity [5, 54]. Even though the diameter effect seems consistent across different conditions, we recommend promoting a variety of large trees of various species as this may further increase the positive effect on biodiversity [7]. Indeed, the succession dynamics and formation rate of microhabitats may vary with tree species [11, 13]. The successional patterns and long-term dynamics of microhabitats remain largely unknown [11] and long-term monitoring at both tree and stand scales are needed to better understand their dynamics and the underlying processes at play [5]. Ultimately, such knowledge will provide robust scientific grounds on which to base biodiversity preservation recommendations for forest managers.

## Acknowledgements

We are in debt to the reserve and forest managers who fed the database and made this study possible. Without their commitment and implication on a daily basis, our results would not have been achievable. In addition, we thank C. Bennemann for her work on GIS data extraction, C. Puverel and R. Andrade for their invaluable help with colour issues and other subtleties of ggplot2, V. Moore for language review and two anonymous reviewers for constructive comments. The data analysed here is issued from the “Protocole de Suivi Des Réserves Forestières” (PSDRF), ONF-RNF.

## Supplementary materials

**Table S1:**
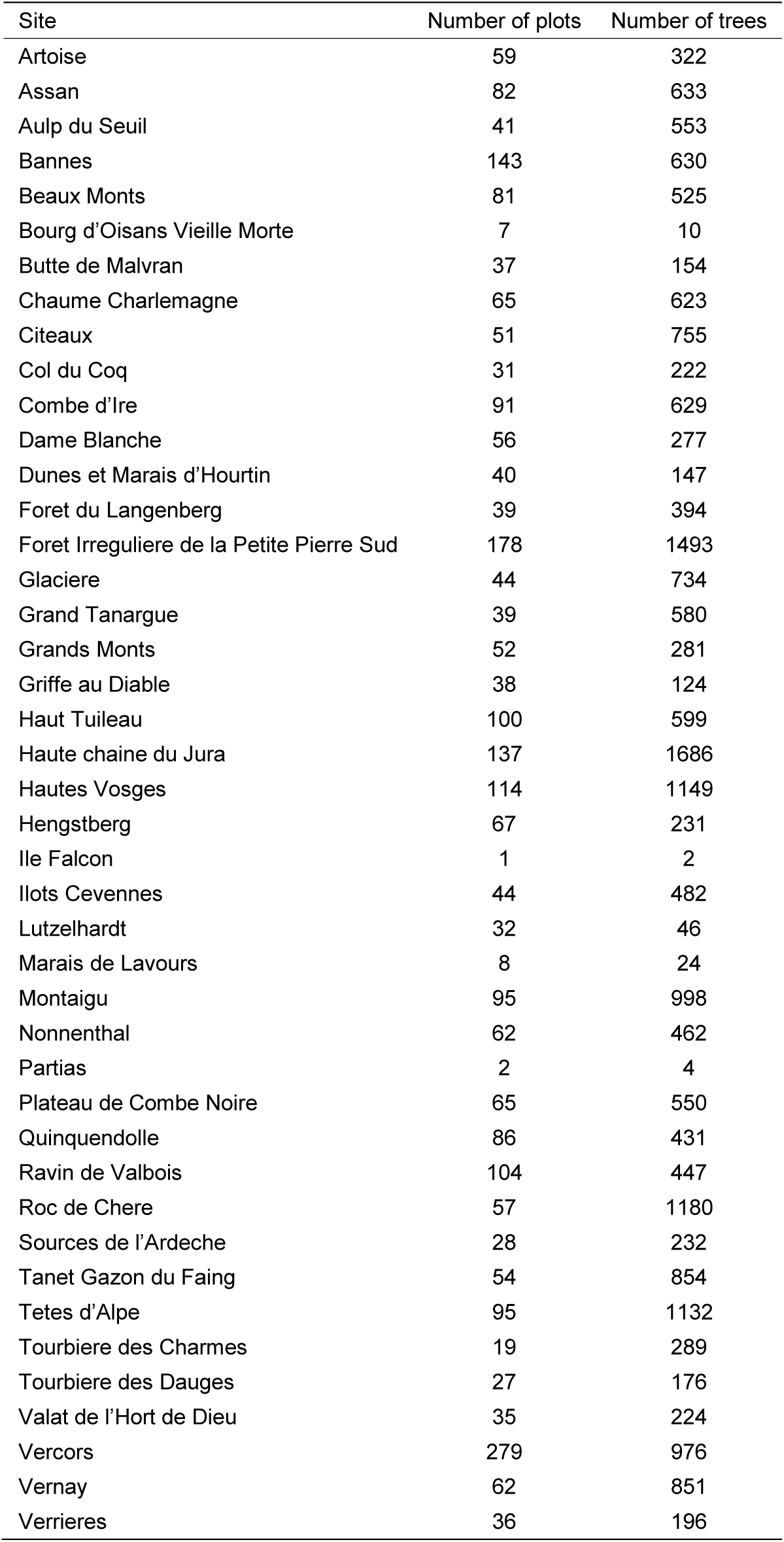
Distribution of plots and trees across the study sites (see map, Figure 1)

**Table S2:** Scaled estimates for number of microhabitat types per tree from a generalised linear mixed model with a Poisson error distribution and plot nested in site as a random effect. DBH: Diameter at Breast Height; SE: standard error of the mean: p = p value; ***p<0.001; **p<0.01; *p<0.05. Beech: Fagus sylvatica; fir: Abies alba; oak: Quercus spp.; pine: Pinus spp.; and spruce: Picea abies.

| Parameter | Estimate | SE | p |  |
| --- | --- | --- | --- | --- |
| Intercept | 0.8198 | 0.0954 | <0.001 | *** |
| DBH | 0.2265 | 0.0359 | <0.001 | *** |
| Fir | -0.3196 | 0.0482 | <0.001 | *** |
| Oak | 0.2060 | 0.0502 | <0.001 | *** |
| Pine | -0.4070 | 0.0838 | <0.001 | *** |
| Spruce | -0.4386 | 0.0558 | <0.001 | *** |
| Living status (Living trees) | -0.3004 | 0.0338 | <0.001 | *** |
| pH | -0.0170 | 0.0506 | 0.7372 | ns |
| Elevation | 0.1136 | 0.0380 | 0.0028 | ** |
| DBH:Fir | -0.0576 | 0.0469 | 0.2193 | ns |
| DBH:Oak | -0.0112 | 0.0474 | 0.8128 | ns |
| DBH:Pine | -0.1282 | 0.0886 | 0.1478 | ns |
| DBH:Spruce | -0.0525 | 0.0541 | 0.3318 | ns |
| DBH: Living status (Living trees) | -0.0098 | 0.0368 | 0.7894 | ns |
| DBH:pH | 0.0460 | 0.0077 | <0.001 | *** |
| Living status (Living trees):pH | -0.0508 | 0.0189 | 0.0072 | ** |
| Fir:pH | -0.0362 | 0.0248 | 0.1445 | ns |
| Oak:pH | 0.0537 | 0.0221 | 0.0153 | * |
| Pine:pH | 0.0976 | 0.0350 | 0.0053 | ** |
| Spruce:pH | -0.0311 | 0.0273 | 0.2553 | ns |
| Fir:Living status (Living trees) | 0.0455 | 0.0491 | 0.3534 | ns |
| Oak:Living status (Living trees) | -0.1354 | 0.0500 | 0.0068 | ** |
| Pine:Living status (Living trees) | 0.0017 | 0.0840 | 0.9837 | ns |
| Spruce:Living status (Living trees) | 0.0708 | 0.0574 | 0.2173 | ns |
| DBH:Fir:Living status (Living trees) | -0.1034 | 0.0491 | 0.0352 | * |
| DBH:Oak:Living status (Living trees) | -0.0160 | 0.0484 | 0.7409 | ns |
| DBH:Pine:Living status (Living trees) | 0.1964 | 0.0954 | 0.0396 | * |
| DBH:Spruce:Living status (Living trees) | -0.1154 | 0.0572 | 0.0435 | * |

**Table S3:** Accumulation levels of microhabitats per tree (number of microhabitats and occurrence) for a Diameter at Breast Height (DBH) increment from 50 cm to 100 cm issued from generalised linear mixed models with Poisson (number) and binomial (occurrence) error distributions. Beech: Fagus sylvatica; fir: Abies alba; oak: Quercus spp.; pine: Pinus spp.; and spruce: Picea abies.

| Microhabitats | Living trees |  |  |  |  | Dead trees |  |  |  |  |
| --- | --- | --- | --- | --- | --- | --- | --- | --- | --- | --- |
|  | Beech | Fir | Oak | Pine | Spruce | Beech | Fir | Oak | Pine | Spruce |
| All | 1.173 | 0.261 | 0.999 | 1.641 | 0.212 | 1.608 | 0.919 | 1.802 | 0.732 | 1.006 |
| Base cavities | 0.176 | 0.064 | 0.052 | 0.427 | 0.026 | 0.345 | 0.063 | 0.061 | -0.004 | 0.181 |
| Trunk cavities | 0.077 | 0.013 | 0.036 | 0.026 | 0.05 | 0.346 | 0.058 | 0.282 | 0.005 | 0.055 |
| Canopy cavities | 0.018 | 0.002 | 0.005 | 0.000 | 0.000 | 0.000 | 0.062 | 0.002 | 0.000 | 0.011 |
| Woodp. cavities | 0.005 | 0.001 | 0.004 | 0.209 | 0.006 | 0.033 | 0.027 | 0.007 | 0.131 | 0.023 |
| Cracks | 0.028 | 0.001 | 0.057 | 0.007 | 0.012 | 0.077 | -0.004 | 0.01 | -0.001 | -0.003 |
| Woodp. feeding holes | 0.032 | 0.005 | 0.024 | 0.002 | 0.004 | 0.417 | 0.305 | 0.411 | 0.022 | 0.233 |
| Rot | 0.011 | 0.003 | 0.033 | 0.042 | 0.001 | 0.234 | 0.007 | 0.267 | 0.039 | -0.019 |
| Injuries | 0.043 | -0.003 | 0.043 | 0.012 | -0.004 | -0.005 | -0.003 | 0.006 | 0.012 | -0.003 |
| Conks of fungi | 0.047 | 0.022 | 0.003 | 0.002 | -0.001 | 0.150 | 0.240 | 0.143 | -0.01 | 0.270 |
| Bark characteristics | 0.004 | 0.002 | 0.013 | 0.003 | 0.000 | 0.021 | 0.09 | 0.121 | 0.014 | -0.028 |
| Moss cover >50% | 0.196 | 0.243 | -0.032 | -0.071 | 0.246 | 0.086 | 0.161 | -0.001 | 0.626 | 0.277 |
| Lichen cover > 50% | 0.063 | 0.097 | 0.017 | 0.174 | -0.045 | -0.017 | 0.02 | 0.005 | -0.027 | 0.026 |
| Ivy cover >50% | 0.001 | 0.002 | 0.058 | -0.002 | 0.006 | 0.001 | 0.000 | 0.003 | 0.011 | 0.000 |
| Small branches | 0.200 | 0.348 | 0.214 | 0.147 | 0.559 | -0.003 | 0.049 | -0.015 | -0.008 | 0.007 |
| Medium branches | 0.498 | 0.36 | 0.526 | 0.324 | 0.019 | 0.015 | 0.010 | 0.007 | -0.007 | 0.013 |
| Large branches | 0.049 | 0.003 | 0.012 | 0.081 | 0.000 | 0.021 | 0.006 | 0.000 | 0.011 | 0.000 |
| Crown skeleton | 0.000 | 0.001 | 0.000 | 0.090 | 0.005 | -0.001 | 0.000 | 0.000 | 0.124 | -0.006 |
| Forks | 0.279 | 0.050 | 0.309 | 0.152 | -0.012 | -0.001 | 0.049 | 0.001 | -0.001 | 0.048 |
| Broken stem | 0.012 | -0.021 | -0.003 | -0.017 | -0.01 | 0.116 | 0.024 | 0.052 | -0.025 | 0.052 |

**Table S4:** Scaled estimates for number of microhabitat types per tree for beech (Fagus sylvatica) and pine (Pinus spp.) from a generalised linear mixed model with a Poisson error distribution and plot nested in site as a random effect. DBH: Diameter at Breast Height; SE: standard error of the mean. p = p value; ***p<0.001; **p<0.01; *p<0.05.

|  | Beech |  |  |  | Pine |  |  |  |
| --- | --- | --- | --- | --- | --- | --- | --- | --- |
|  | Estimate | SE | p |  | Estimate | SE | p |  |
| Intercept | 0.759 | 0.097 | <0.001 | *** | 0.285 | 0.186 | 0.126 | ns |
| DBH | 0.147 | 0.034 | <0.001 | *** | 0.101 | 0.046 | 0.030 | * |
| Living status (Living trees) | -0.270 | 0.033 | <0.001 | *** | -0.441 | 0.069 | <0.001 | *** |
| pH | 0.032 | 0.064 | 0.614 | ns | 0.173 | 0.190 | 0.363 | ns |
| Elevation | -0.030 | 0.060 | 0.617 | ns | -0.250 | 0.168 | 0.137 | ns |
| DBH:Living status (Living trees) | 0.030 | 0.034 | 0.375 | ns | 0.124 | 0.051 | 0.016 | * |
| DBH:pH | 0.045 | 0.010 | <0.001 | *** | 0.169 | 0.067 | 0.012 | * |
| DBH:Elevation | -0.007 | 0.009 | 0.477 | ns | -0.105 | 0.064 | 0.097 | (*) |
| Living status (Living trees):pH | -0.078 | 0.033 | 0.017 | * | -0.171 | 0.155 | 0.269 | ns |
| Living status (Living trees):Elevation | 0.120 | 0.036 | 0.001 | ** | 0.302 | 0.166 | 0.068 | (*) |

**Table S5:**
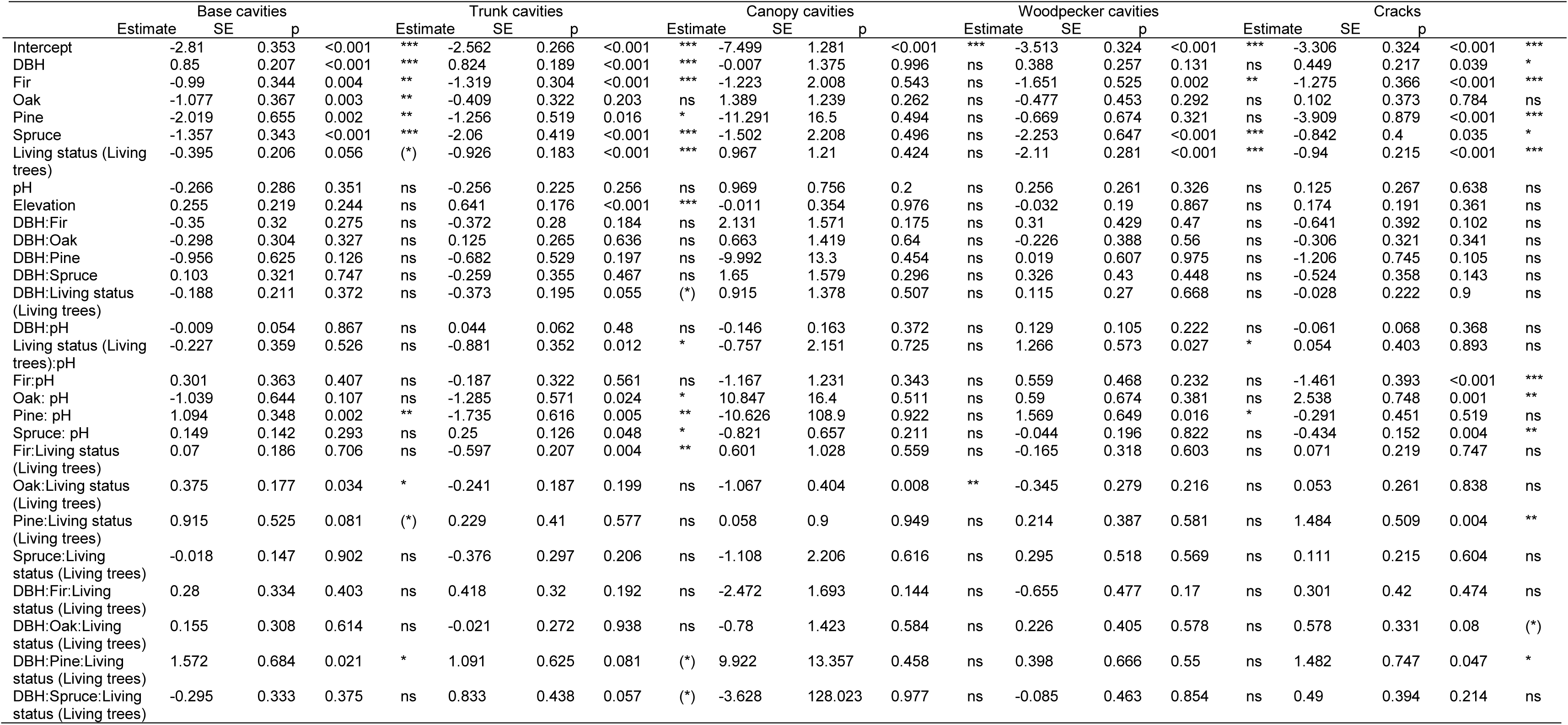
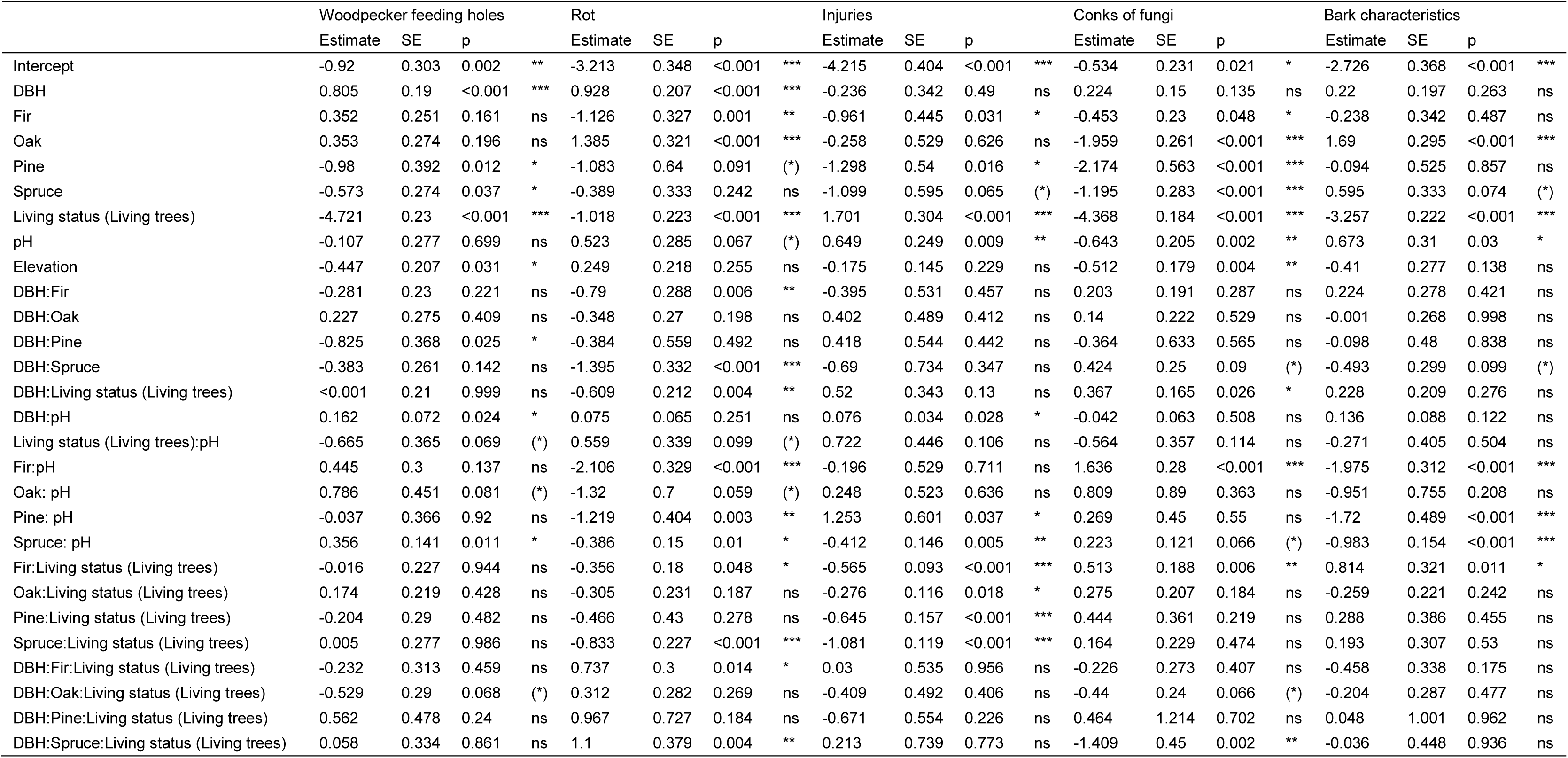
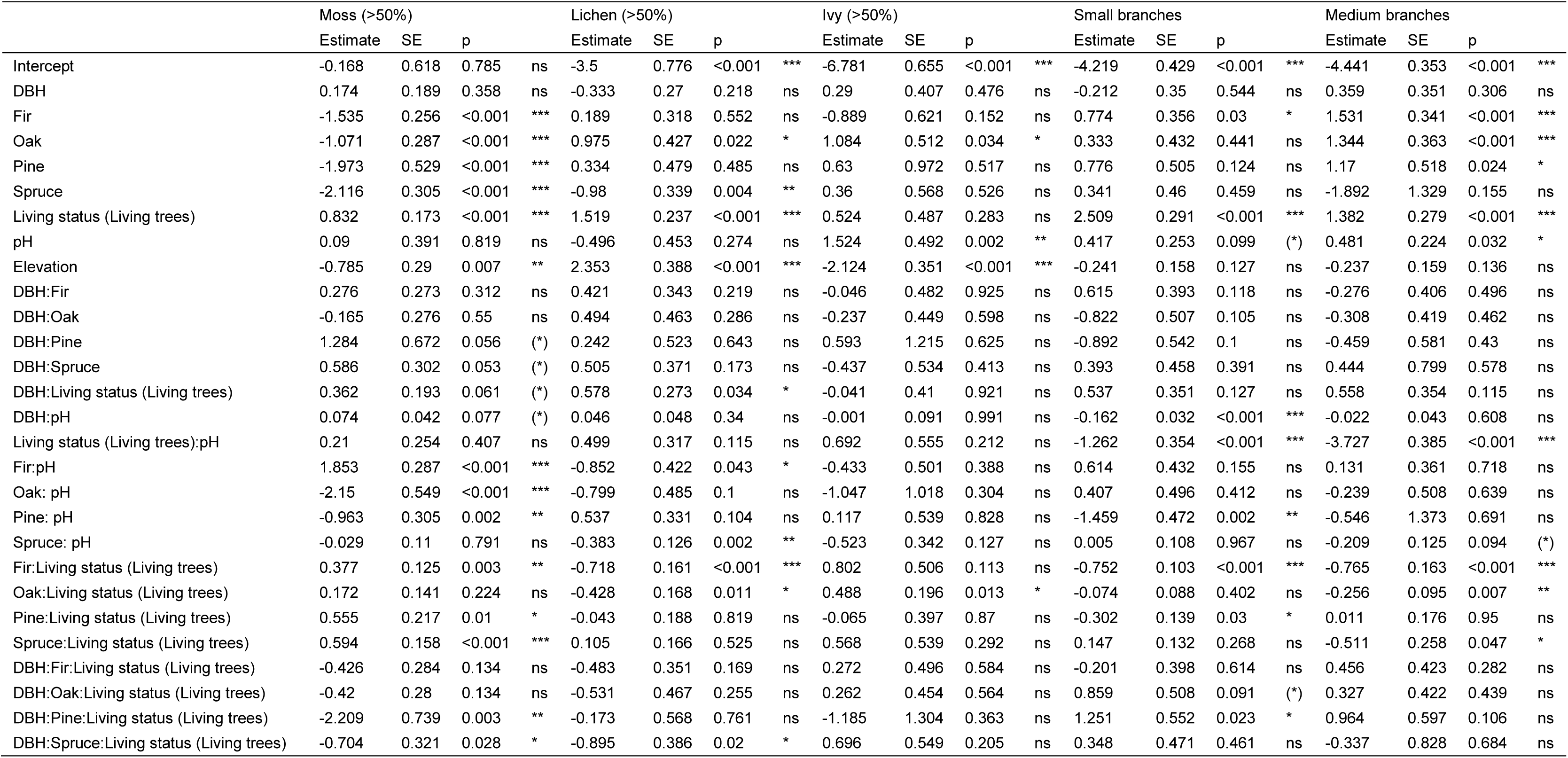
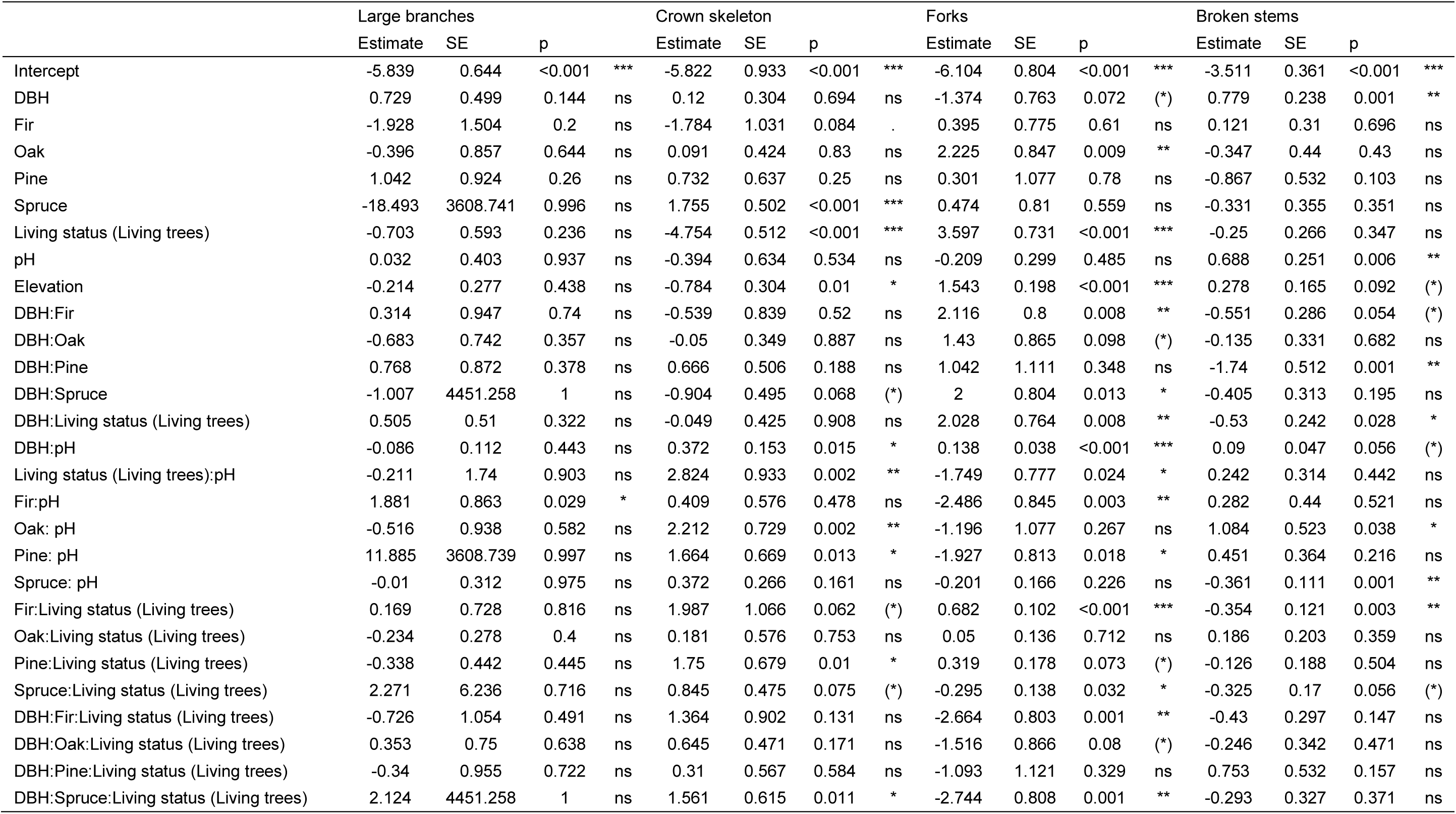
Scaled estimates for occurrence of microhabitat types per tree from a generalised linear mixed model with a binomial error distribution and plot nested in site as a random effect. DBH: Diameter at Breast Height; SE: standard error of the mean. Beech: Fagus sylvatica; fir: Abies alba; oak: Quercus spp.; pine: Pinus spp.; and spruce: Picea abies. p = p value; ***p<0.001; **p<0.01; *p<0.05.

**Figure S1:**
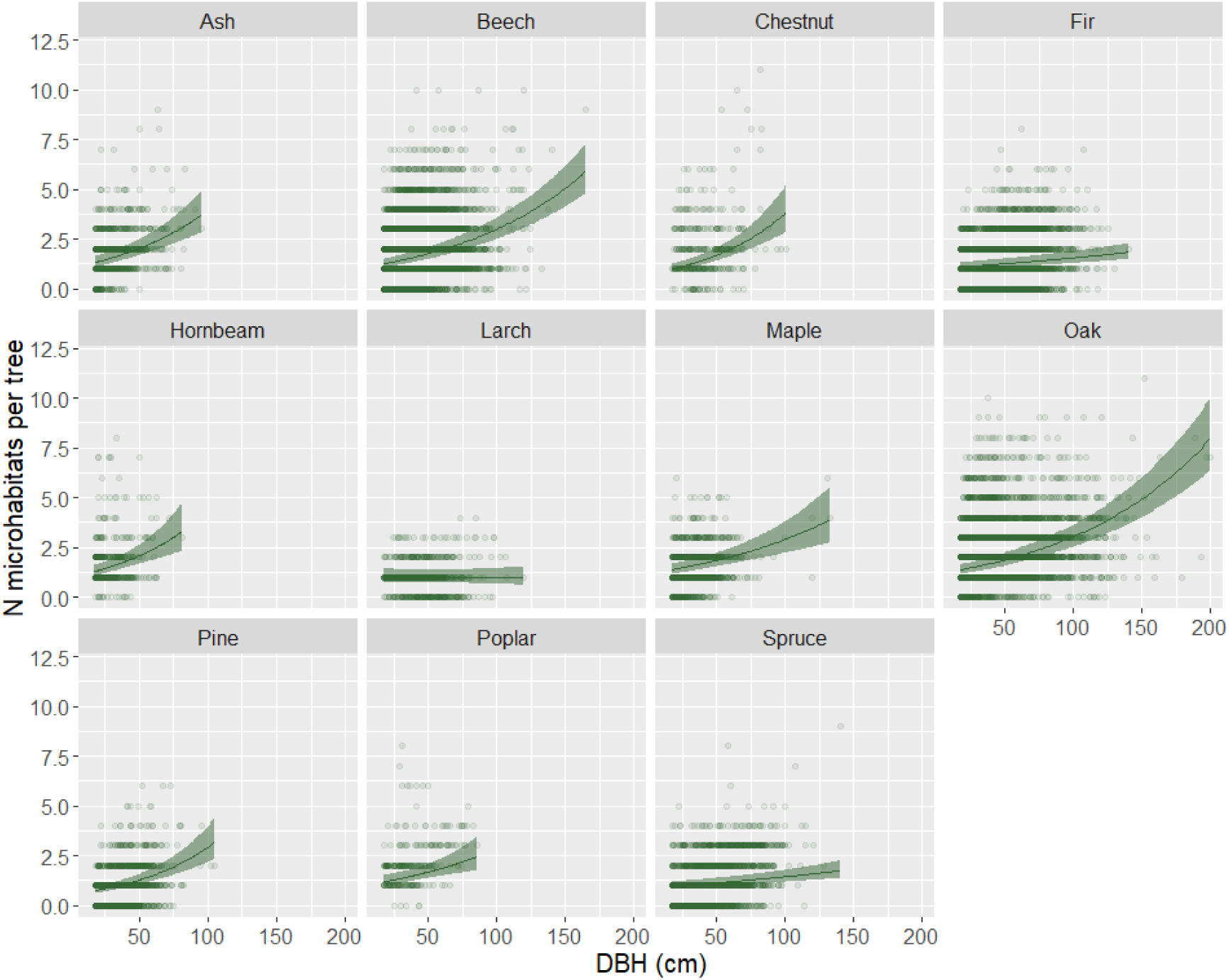
Relationship between total number of microhabitats (N microhabitats per tree) and Diameter at Breast Height (DBH, cm) by genus for living trees only. Lines represent estimates from generalized mixed effect models with a Poisson error distribution. Ribbons show the 95% confidence interval of the mean. For this representation, pH and elevation were held constant. Ash: Fraxinus excelsior; beech: Fagus sylvatica; chestnut: Castanea sativa; fir: Abies alba; hornbeam: Carpinus betulus; larch: Larix decidua; maple: Acer spp., oak: Quercus spp.; pine: Pinus spp.; poplar: Populus spp.; and spruce: Picea abies.

**Figure S2:**
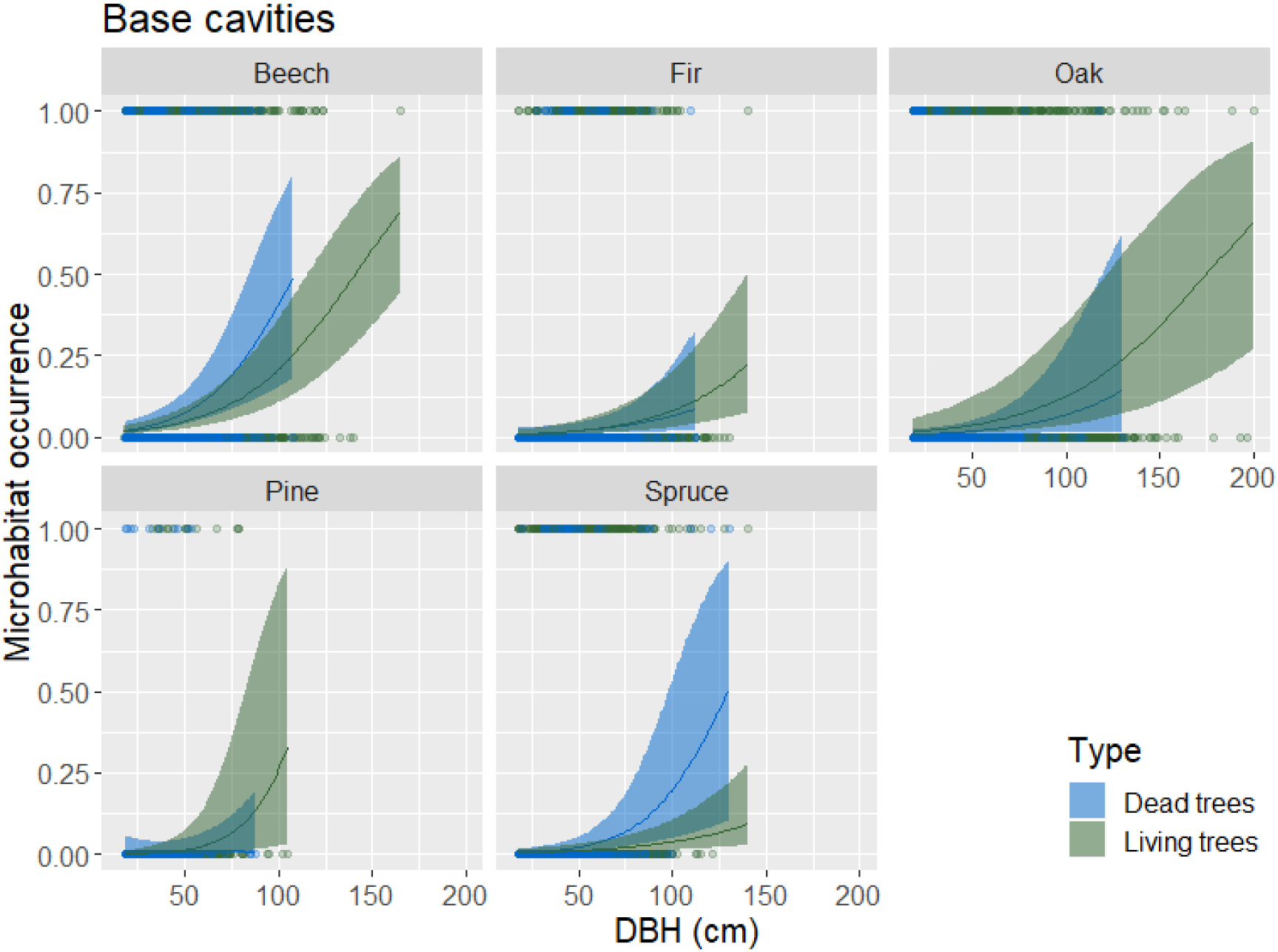
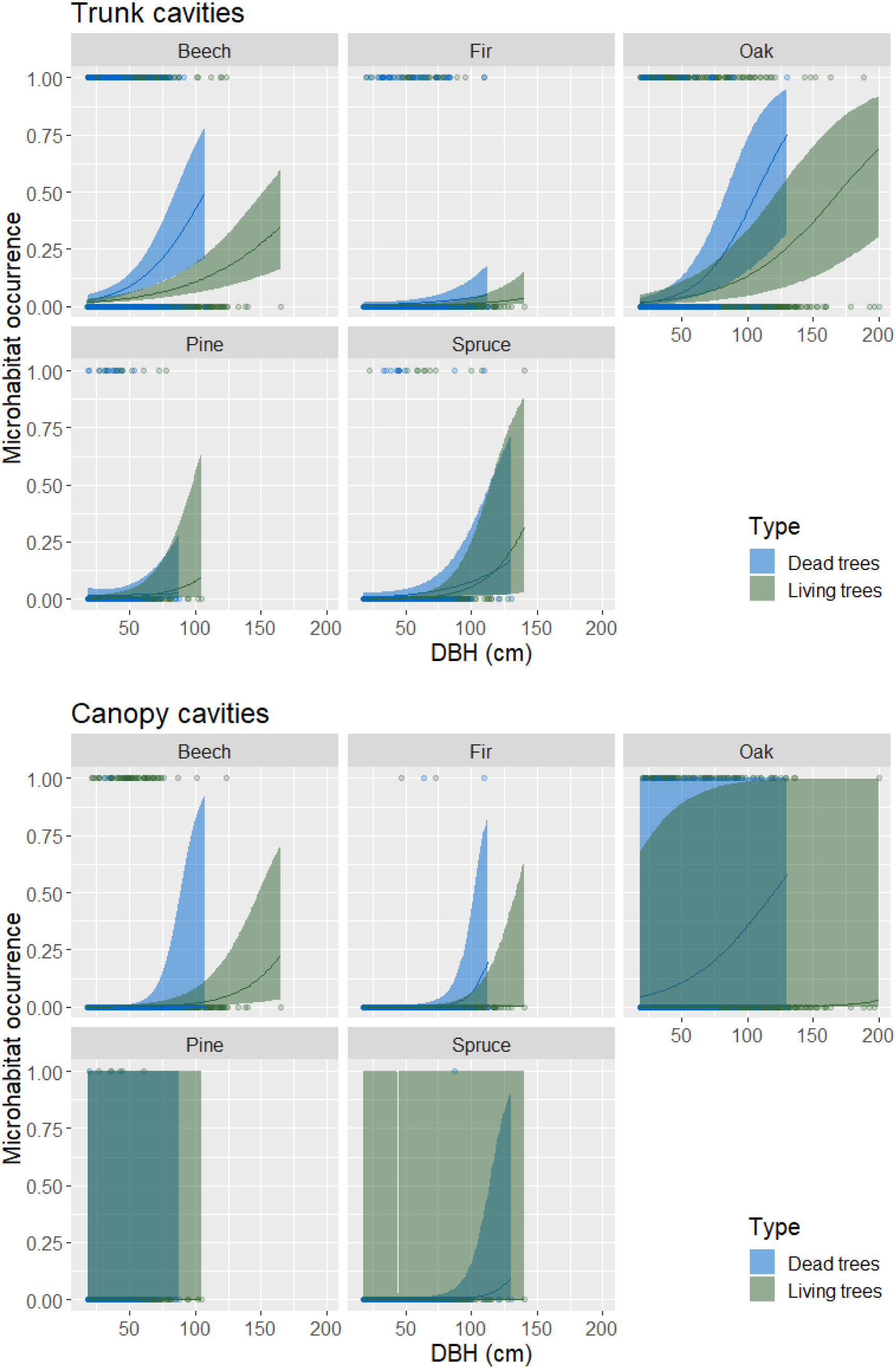
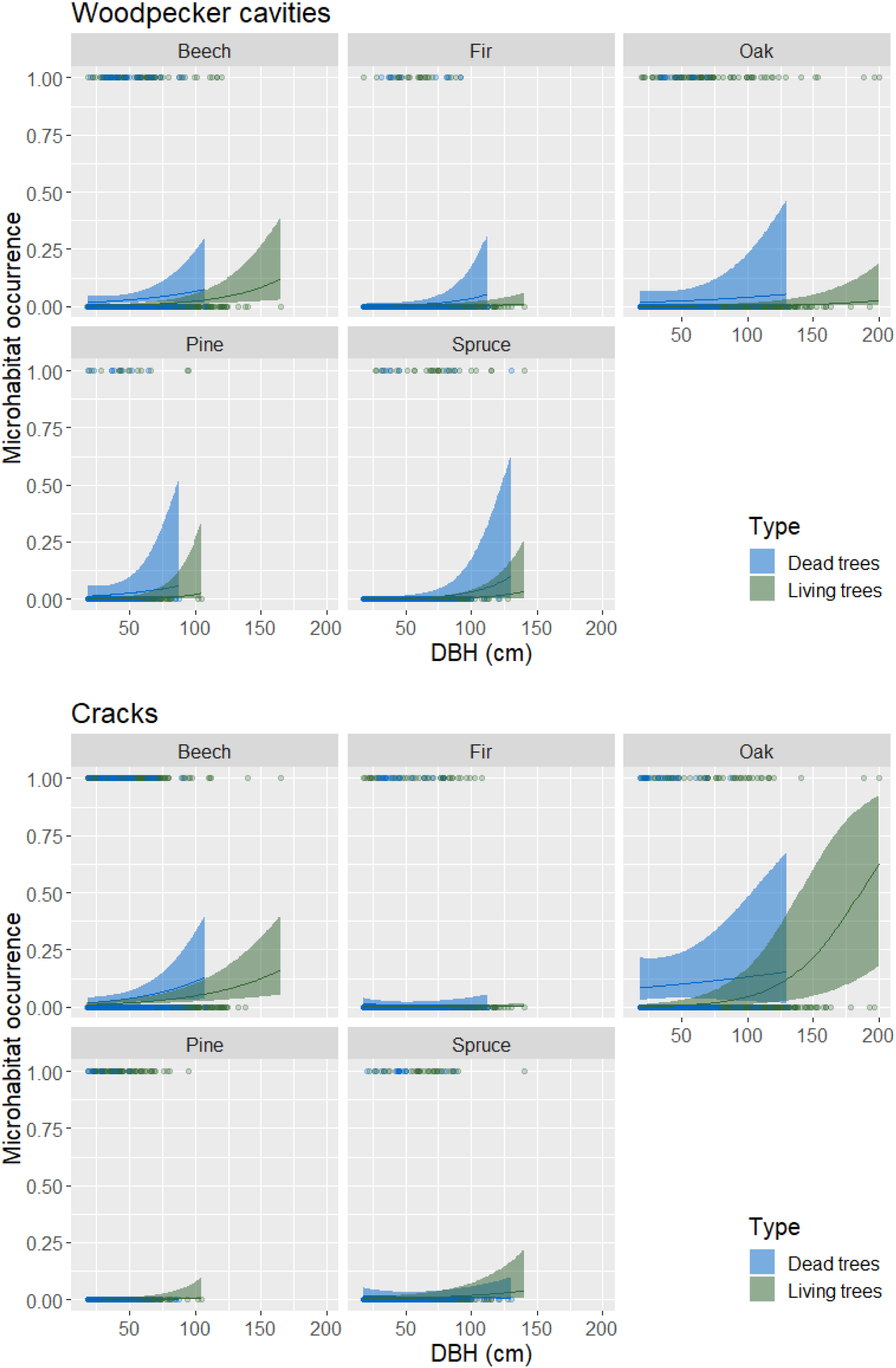
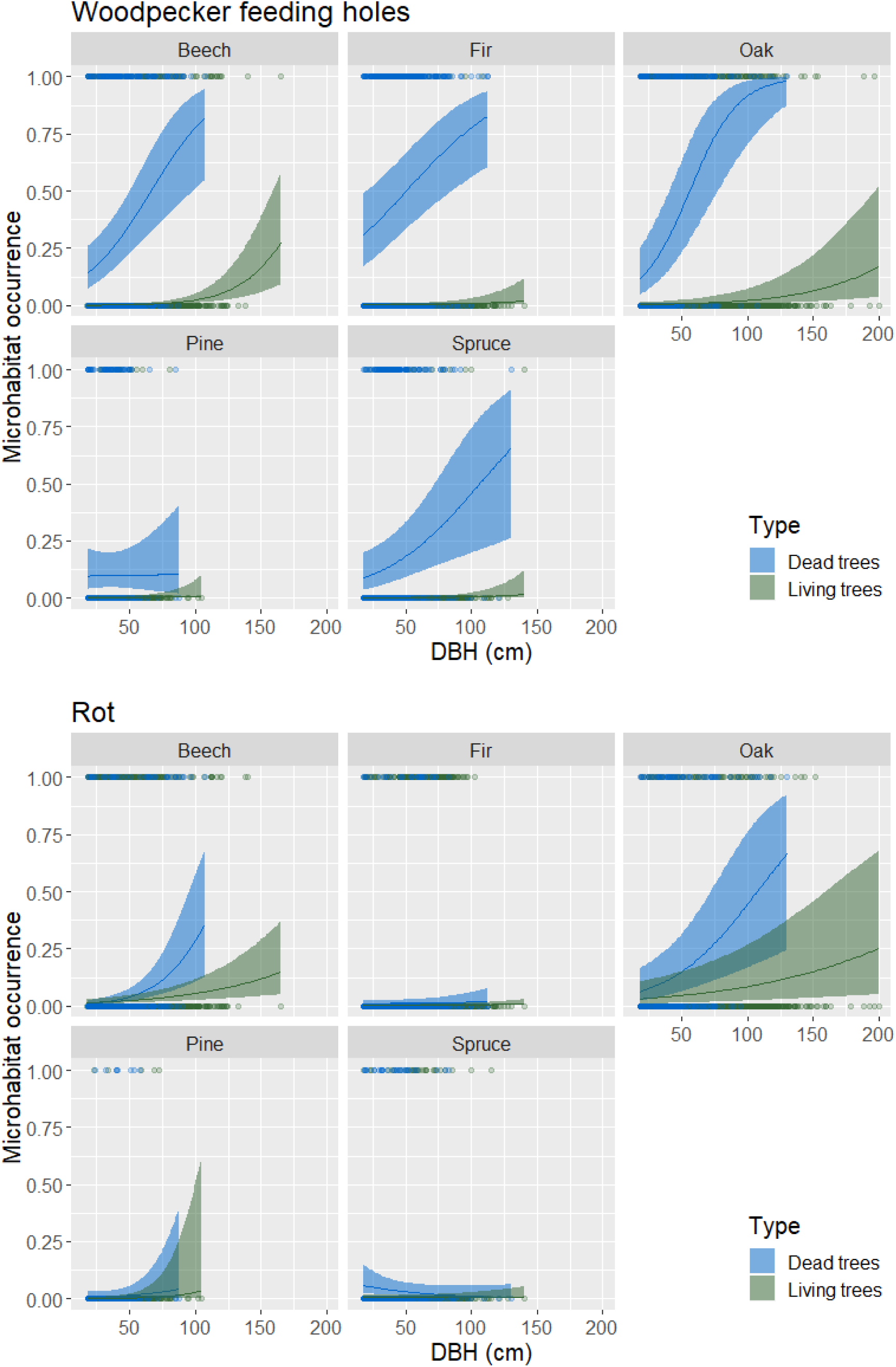
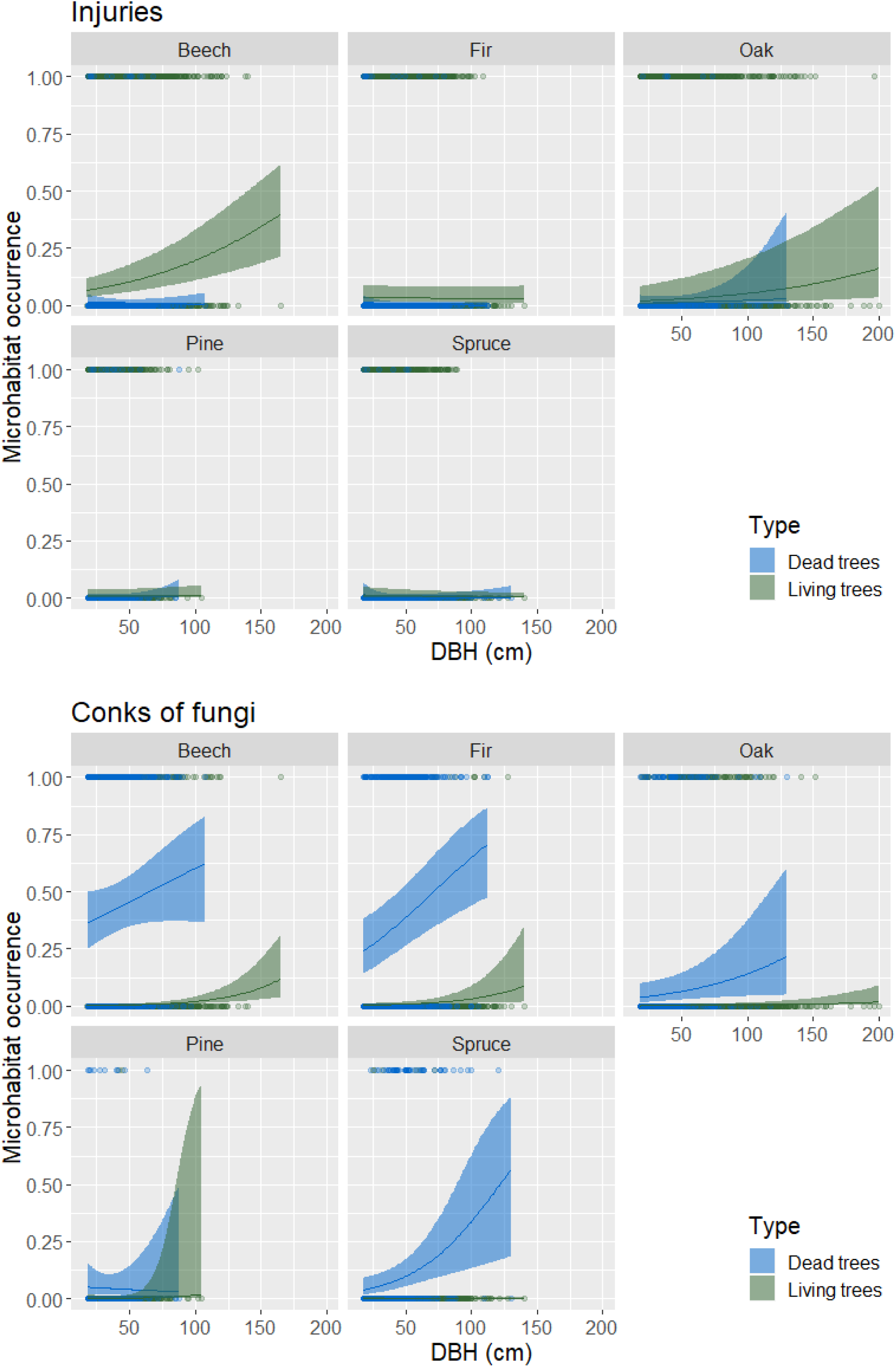
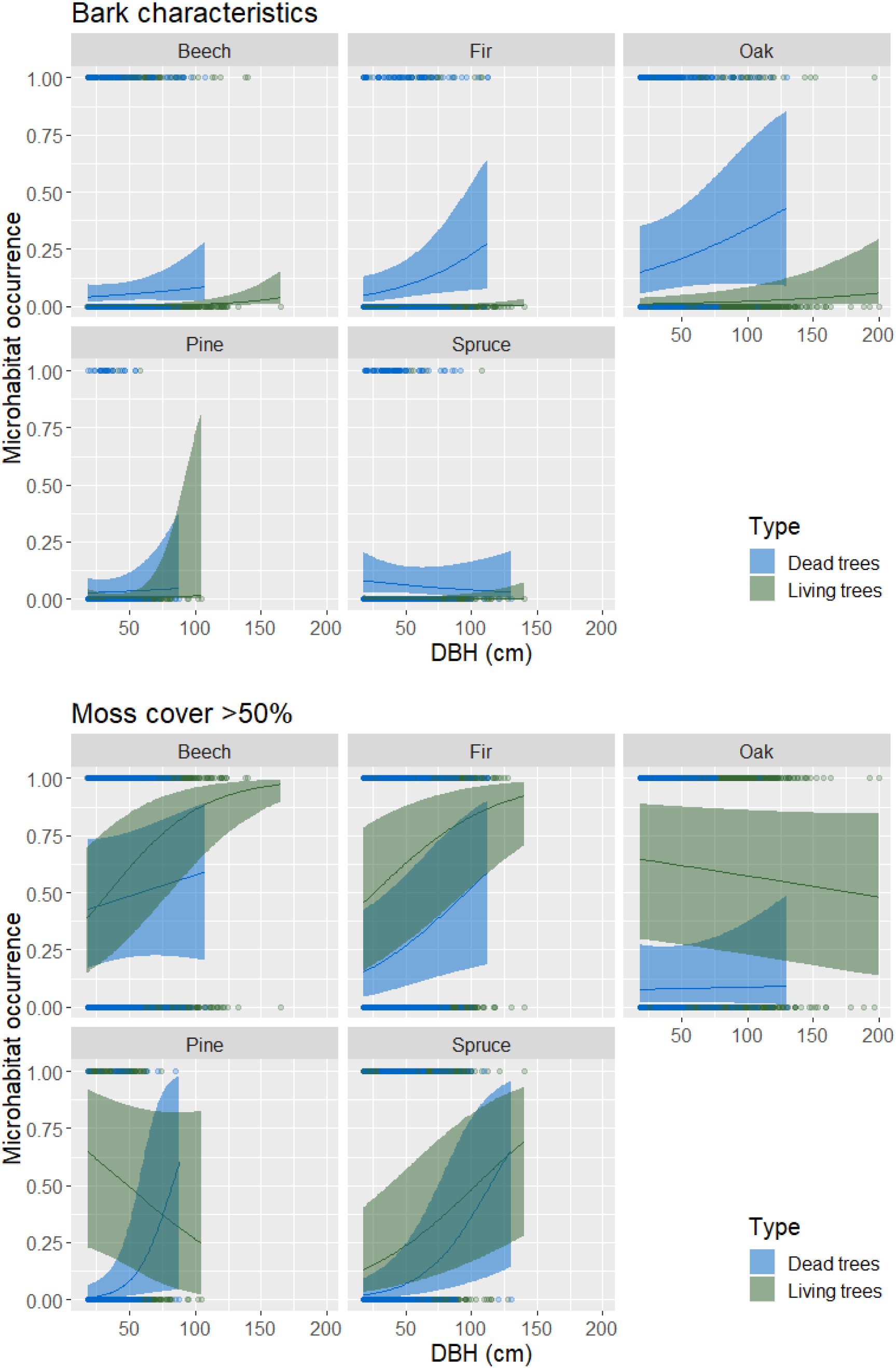
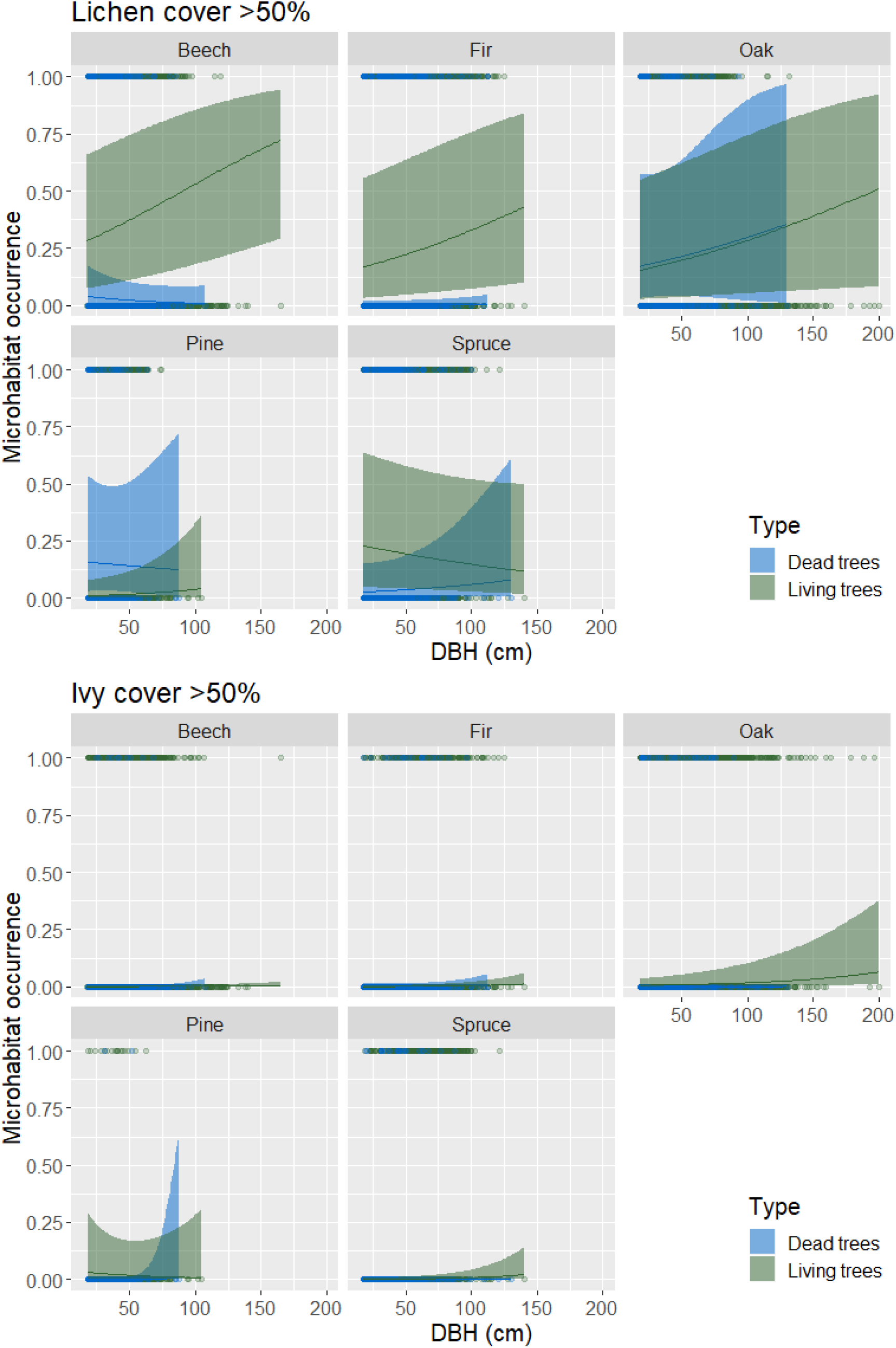
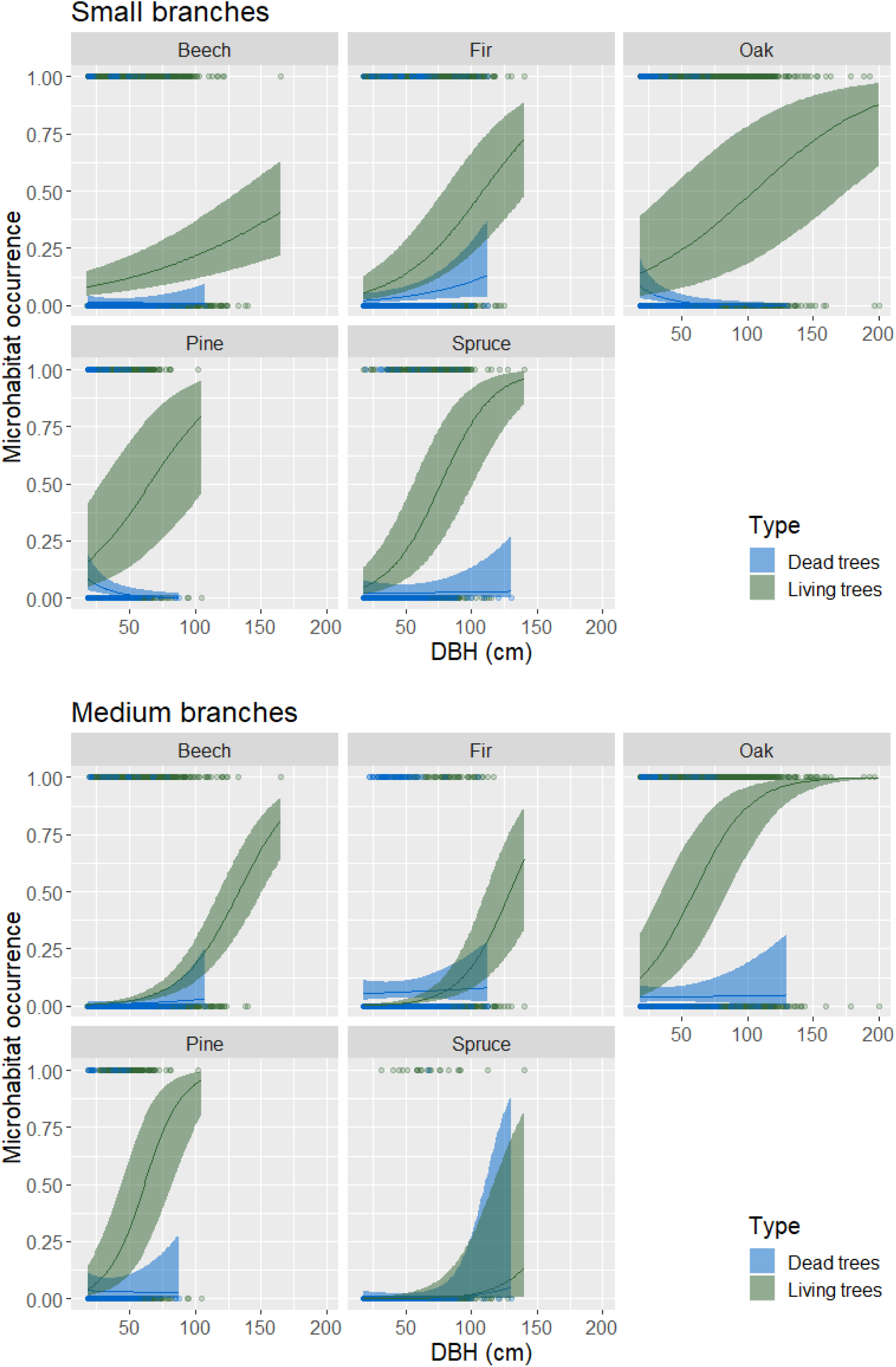
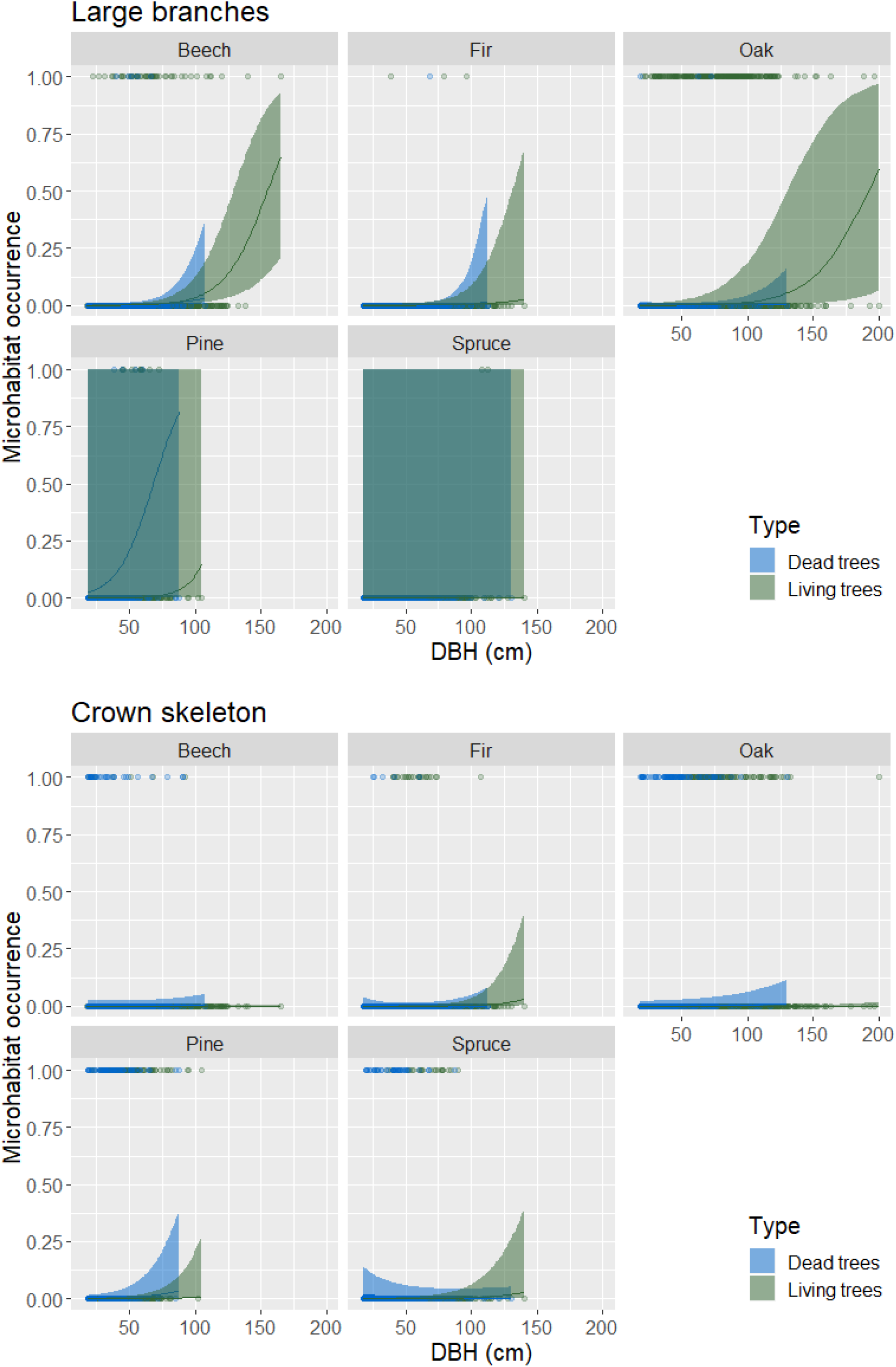
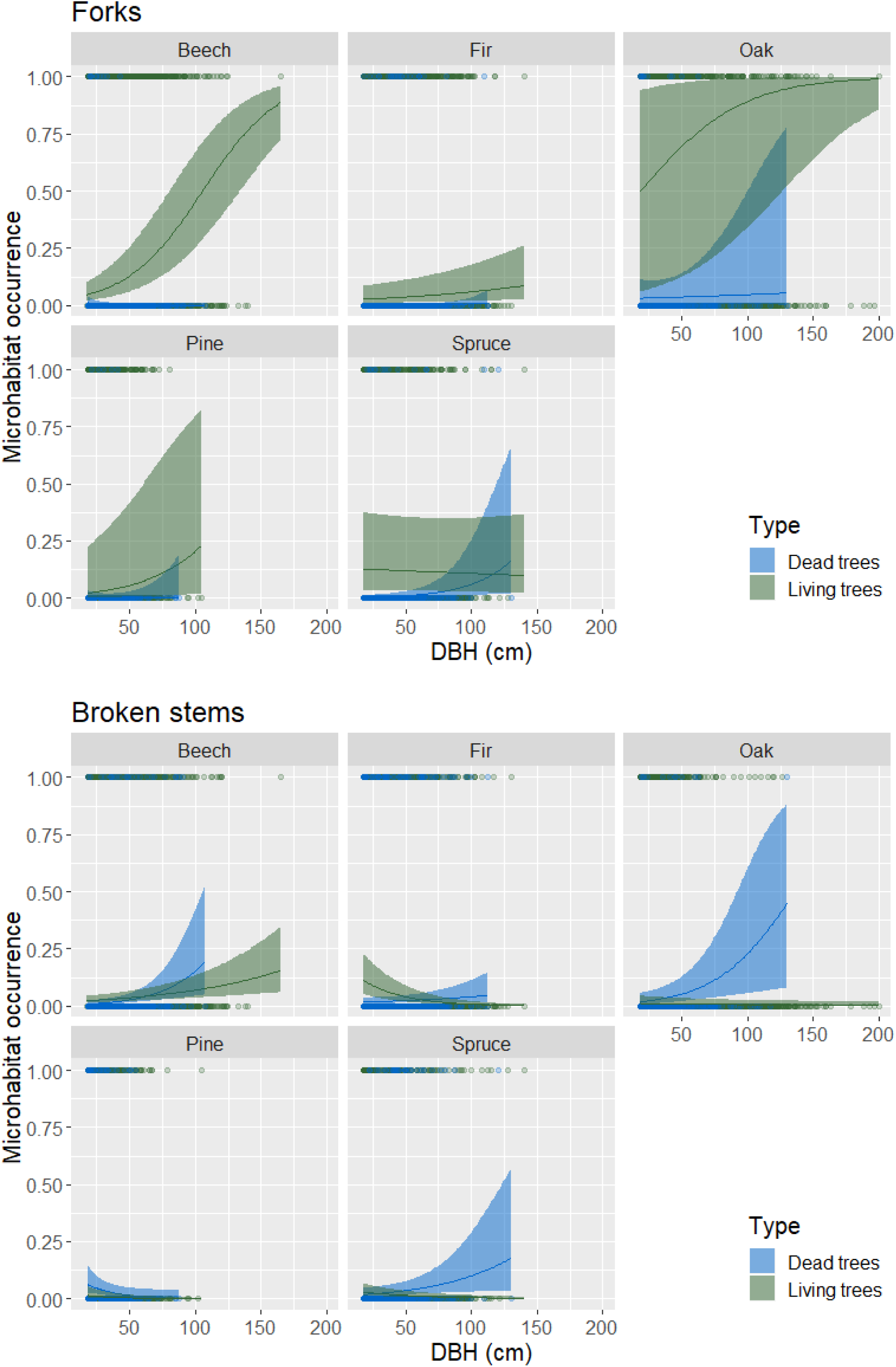
Relationship between occurrence of microhabitats per tree and Diameter at Breast Height (DBH, cm) by species and living status (living vs. dead standing trees). Lines represent estimates from generalized mixed effect models with a binomial error distribution. Ribbons show the 95% confidence interval of the mean. For the representation, pH and elevation were held constant. Beech: Fagus sylvatica; fir: Abies alba; oak: Quercus spp.; pine: Pinus spp.; and spruce: Picea abies.

## References cited

1. Hunter ML, Acuña V, Bauer DM, Bell KP, Calhoun AJK, Felipe-Lucia MR, et al. Conserving small natural features with large ecological roles: A synthetic overview. Biol Conserv. 2017;211:88–95. doi: 10.1016/j.biocon.2016.12.020.

2. Bauer DM, Bell KP, Nelson EJ, Calhoun AJK. Managing small natural features: A synthesis of economic issues and emergent opportunities. Biol Conserv. 2017;211:80–7. doi: 10.1016/j.biocon.2017.01.001.

3. Lindenmayer DB. Conserving large old trees as small natural features. Biol Conserv. 2017;211:51–9. doi: 10.1016/j.biocon.2016.11.012.

4. Lindenmayer DB, Laurance WF, Franklin JF. Global decline in large old trees. Science. 2012;335:1305–6. doi: 10.1126/science.1231070.

5. Larrieu L, Paillet Y, Winter S, Bütler R, Kraus D, Krumm F, et al. Tree related microhabitats in temperate and Mediterranean European forests: A hierarchical typology for inventory standardization. Ecol Indic. 2018;84(April 2017):194–207. doi: 10.1016/j.ecolind.2017.08.051.

6. Paillet Y, Archaux F, Boulanger V, Debaive N, Fuhr M, Gilg O, et al. Snags and large trees drive higher tree microhabitat densities in strict forest reserves. For Ecol Manag. 2017;389:176–86. doi: 10.1016/j.foreco.2016.12.014.

7. Paillet Y, Archaux F, du Puy S, Bouget C, Boulanger V, Debaive N, et al. The indicator side of tree microhabitats: a multi-taxon approach based on bats, birds and saproxylic beetles. J Appl Ecol. 2018;(April):0–1. doi: 10.1111/1365-2664.13181.

8. Regnery B, Couvet D, Kubarek L, Julien JF, Kerbiriou C. Tree microhabitats as indicators of bird and bat communities in Mediterranean forests. Ecol Indic. 2013;34:221–30.

9. Bütler R, Lachat T, Larrieu L, Paillet Y. Habitat trees: key elements for forest biodiversity. In: Kraus D, Krumm F, editors. Integrative approaches as an opportunity fir the conservation of forest biodiversity. Freiburg, DEU: European Forest Institute; 2013. p. 84–91.

10. Stokland JN, Siitonen J, Jonsson BG. Biodiversity in dead wood. Cambridge, UK: University Press; 2012.

11. Courbaud B, Pupin C, Letort A, Cabanettes A, Larrieu L, Börger L. Modelling the probability of microhabitat formation on trees using cross-sectional data. Meth Ecol Evol. 2017;8(10):1347–59. doi: 10.1111/2041-210x.12773.

12. Larrieu L, Cabanettes A. Species, live status, and diameter are important tree features for diversity and abundance of tree microhabitats in subnatural montane beech-fir forests. Can J For Res. 2012;42(8):1433–45.

13. Vuidot A, Paillet Y, Archaux F, Gosselin F. Influence of tree characteristics and forest management on tree microhabitats. Biol Conserv. 2011;144(1):441–50. doi: 10.1016/j.biocon.2010.09.030.

14. Winter S, Höfler J, Michel AK, Böck A, Ankerst DP. Association of tree and plot characteristics with microhabitat formation in European beech and Douglas-fir forests. European Journal of Forest Research. 2015;134(2):335–47. doi: 10.1007/s10342-014-0855-x.

15. Siitonen J. Microhabitats. In: Stokland JN, Siitonen J, Jonsson BG, editors. Biodiversity in dead wood. New York, USA: Cambridge University Press; 2012. p. 150–82.

16. Regnery B, Paillet Y, Couvet D, Kerbiriou C. Which factors influence the occurrence and density of tree microhabitats in Mediterranean oak forests? For Ecol Manag. 2013;295:118–25.

17. Großmann J, Schultze J, Bauhus J, Pyttel P. Predictors of Microhabitat Frequency and Diversity in Mixed Mountain Forests in South-Western Germany. Forests. 2018;9(3):104. doi: 10.3390/f9030104.

18. Winter S, Möller GC. Microhabitats in lowland beech forests as monitoring tool for nature conservation. For Ecol Manag. 2008;255(3-4):1251–61.

19. Kraus D, Schuck A, Bebi P, Blaschke M, Bütler R, Flade M, et al. Spatially explicit database of tree related microhabitats (TreMs). Version 1.2. ed. GBIF.org: Integrate+ project, Institut National de la Recherche Agronomique (INRA); 2017.

20. Paillet Y, Pernot C, Boulanger V, Debaive N, Fuhr M, Gilg O, et al. Quantifying the recovery of old-growth attributes in forest reserves: A first reference for France. For Ecol Manag. 2015;346:51–64. doi: 10.1016/j.foreco.2015.02.037.

21. Kraus D, Bütler R, Krumm F, Lachat T, Larrieu L, Mergner U, et al. Catalogue of tree microhabitats – Reference field list. Integrate+ Technical Paper 16p. 2016:16-. doi: 10.13140/RG.2.1.1500.6483.

22. Paillet Y, Coutadeur P, Vuidot A, Archaux F, Gosselin F. Strong observer effect on tree microhabitats inventories: A case study in a French lowland forest. Ecol Indic. 2015;49:14–23. doi: 10.1016/j.ecolind.2014.08.023.

23. Fick SE, Hijmans RJ. WorldClim 2: new 1-km spatial resolution climate surfaces for global land areas. International Journal of Climatology. 2017;37(12):4302–15. doi: 10.1002/joc.5086.

24. Gégout J-C. Validation des bio-indicateurs floristiques pour une surveillance de l’état nutritionnel des sols forestiers français à partir des données de l’Inventaire forestier national - Rapport final,. Convention ADEME/IFN/ENGREF n° 0562C0029, 2008,.

25. Zuur AF, Ieno EN, Elphick CS. A protocol for data exploration to avoid common statistical problems. Meth Ecol Evol. 2010;1(1):3–14. doi: 10.1111/j.2041-210X.2009.00001.x. PubMed PMID: WOS:000288913700002.

26. Magnusson A, Skaug HJ, Nielsen A, Berg CW, Kristensen K, Maechler M, et al. glmmTMB: Generalized Linear Mixed Models using Template Model Builder. R package version 0.1.3. ed2017.

27. Lenth R. emmeans: Estimated Marginal Means, aka Least-Squares Means. R package version 1.1. ed2018.

28. Sellers KF, Morris DS. Underdispersion models: Models that are “under the radar”. Communications in Statistics - Theory and Methods. 2017;46(24):12075–86. doi: 10.1080/03610926.2017.1291976.

29. Barbier S, Chevalier R, Loussot P, Bergès L, Gosselin F. Improving biodiversity indicators of sustainable forest management: Tree genus abundance rather than tree genus richness and dominance for understory vegetation in French lowland oak hornbeam forests. For Ecol Manag. 2009;258:S176–S86. doi: 10.1016/j.foreco.2009.09.004. PubMed PMID: ISI:000273140300019.

30. R Core Team. R: A language and environment for statistical computing. Vienna, Austria: R Foundation for Statistical Computing; 2017.

31. Cockle KL, Martin K, Wesołowski T. Woodpeckers, decay, and the future of cavity-nesting vertebrate communities worldwide. Frontiers in Ecology and the Environment. 2011;9(7):377–82. doi: 10.1890/110013.

32. Jackson JA, Jackson BJS. Ecological relationships between fungi and woodpecker cavity sites. Condor. 2004;106(1):37–49.

33. Laiolo P, Rolando A, Valsania V. Responses of birds to the natural re-establishment of wilderness in montane beechwoods of North-western Italy. Acta Oecol. 2004;25(1-2):129–36. doi: 10.1016/j.actao.2003.12.003.

34. Zarnowitz JE, Manuwal DA. The effects of forest management on cavity-nesting birds in northwestern Washington. J Wildl Manag. 1985;49(1):255–63.

35. Gouix N, Brustel H. Emergence trap, a new method to survey Limoniscus violaceus (Coleoptera: Elateridae) from hollow trees. Biodivers Conserv. 2012;21(2):421–36. doi: 10.1007/s10531-011-0190-1.

36. Mežaka A, Brumelis G, Piterans A. Tree and stand-scale factors affecting richness and composition of epiphytic bryophytes and lichens in deciduous woodland key habitats. Biodivers Conserv. 2012;21(12):3221–41. doi: 10.1007/s10531-012-0361-8.

37. Ódor P, Király I, Tinya F, Bortignon F, Nascimbene J. Reprint of: Patterns and drivers of species composition of epiphytic bryophytes and lichens in managed temperate forests. For Ecol Manag. 2014;321:42–51. doi: 10.1016/j.foreco.2014.01.035.

38. Johann F, Schaich H. Land ownership affects diversity and abundance of tree microhabitats in deciduous temperate forests. For Ecol Manag. 2016;380:70–81. doi: 10.1016/j.foreco.2016.08.037.

39. Bobiec A. Living stands and dead wood in the Białowieża forest: Suggestions for restoration management. For Ecol Manag. 2002;165(1-3):125–40.

40. Bouget C, Brin A, Brustel H. Exploring the “last biotic frontier”: Are temperate forest canopies special for saproxylic beetles? For Ecol Manag. 2011;261(2):211–20.

41. Remm J, Lõhmus A. Tree cavities in forests - The broad distribution pattern of a keystone structure for biodiversity. For Ecol Manag. 2011;262(4):579–85.

42. Tillon L, Bresso K, Aulagnier S. Tree selection by roosting bats in a European temperate lowland sub-Atlantic forest. Mammalia. 2016;80(3):271–9. doi: 10.1515/mammalia-2014-0095.

43. Scheffers BR, Edwards DP, Diesmos A, Williams SE, Evans TA. Microhabitats reduce animal’s exposure to climate extremes. Global Change Biology. 2014;20(2):495–503. doi: 10.1111/gcb.12439.

44. Edworthy AB, Martin K. Persistence of tree cavities used by cavity-nesting vertebrates declines in harvested forests. J Wildl Manag. 2013;77(4):770–6. doi: 10.1002/jwmg.526.

45. Wesołowski T. “Lifespan” of woodpecker-made holes in a primeval temperate forest: A thirty year study. For Ecol Manag. 2011;262(9):1846–52. doi: 10.1016/j.foreco.2011.08.001.

46. Halme P, Kotiaho JS. The importance of timing and number of surveys in fungal biodiversity research. Biodivers Conserv. 2012;21(1):205–19.

47. Aakala T, Kuuluvainen T, Gauthier S, De Grandpré L. Standing dead trees and their decay-class dynamics in the northeastern boreal old-growth forests of Quebec. For Ecol Manag. 2008;255(3-4):410–20. doi: 10.1016/j.foreco.2007.09.008.

48. Ding Y, Liu G, Zang R, Zhang J, Lu X, Huang J. Distribution of vascular epiphytes along a tropical elevational gradient: Disentangling abiotic and biotic determinants. Scientific Reports. 2016;6. doi: 10.1038/srep19706.

49. Parviainen J. Virgin and natural forests in the temperate zone of Europe. For Snow Landsc Res. 2005;79(1-2):9–18.

50. Christensen M, Hahn K, Mountford EP, Odor P, Standovar T, Rozenbergar D, et al. Dead wood in European beech (Fagus sylvatica) forest reserves. For Ecol Manag. 2005;210(1-3):267–82. PubMed PMID: ISI:000229165500020.

51. Sabatini FM, Burrascano S, Keeton WS, Levers C, Lindner M, Pötzschner F, et al. Where are Europe’s last primary forests? Divers Distrib. 2018. doi: 10.1111/ddi.12778.

52. Kozák D, Mikoláš M, Svitok M, Bače R, Paillet Y, Larrieu L, et al. Profiles of tree-related microhabitats in European primary beech-dominated forests. For Ecol Manag. 2018;429:363–74.

53. Bauhus J, Puettmann K, Messier C. Silviculture for old-growth attributes. For Ecol Manag. 2009;258(4):525–37. doi: 10.1016/j.foreco.2009.01.053.

54. Gossner MM, Getzin S, Lange M, Pašalić E, Türke M, Wiegand K, et al. The importance of heterogeneity revisited from a multiscale and multitaxa approach. Biol Conserv. 2013;166:212–20. doi: 10.1016/j.biocon.2013.06.033.

